# *SELF-PRUNING 5G* interplays with age and gibberellin pathways downstream of *SlCOL1* to orchestrate photoperiodic tomato flowering

**DOI:** 10.64898/2026.08.04.742724

**Authors:** Mateus Henrique Vicente, Gloria Serrano-Bueno, Kailash Pandey, Alan Cesar Faustinoni Fernandes, Flaviani Gabriela Pierdona, Rafael Rubino, Yajahaira Nevenka Carbajal Gonzales, Ramon Gabriel, Carolina Camacho Fernandez, Manuela Rodríguez Delgado, Lilian Ellen Pino, Pedro de los Reyes, Fernando Baile, Lázaro Eustaquio Pereira Peres, Myriam Calonje, Marian Bemer, Federico Valverde, Fabio Tebaldi Silveira Nogueira

**Affiliations:** Laboratory of Molecular Genetics of Plant Development, Escola Superior de Agricultura “Luiz de Queiroz” (ESALQ), University of São Paulo (USP), Piracicaba, São Paulo, Brazil; Plant Development Group - Institute for Plant Biochemistry and Photosynthesis, Consejo Superior de Investigaciones Científicas, Universidad de Sevilla, Seville, Spain; Department of Plant Biochemistry and Molecular Biology, Universidad de Sevilla, Seville, Spain; Biosystematics group, Wageningen University & Research, 6708 PB Wageningen, the Netherlands; Plant Development Group - Institute for Plant Biochemistry and Photosynthesis, Consejo Superior de Investigaciones Científicas-Universidad de Sevilla 41092, Seville, Spain; Laboratory of Plant Developmental Genetics, Escola Superior de Agricultura “Luiz de Queiroz” (ESALQ), University of São Paulo (USP), Piracicaba, São Paulo, Brazil

**Keywords:** Photoperiod, *CONSTANS*, SP5G, Flowering, Gibberellin, miR156

## Abstract

Tomato (*Solanum lycopersicum*) is classified as a day-neutral plant, whereas its wild relatives exhibit delayed flowering under long day (LD) conditions due to higher activity of the *SELF-PRUNING 5G* (*SP5G*). In *Arabidopsis thaliana*, CONSTANS (CO) activates *FLOWERING LOCUS T* (*FT*), a homolog of *SP5G*, but whether and how CO integrates with *SP5G* in tomato flowering was unclear. Here, we demonstrate that *SlCOL1* (the tomato CO homolog) delays flowering by directly activating *SP5G* in a *PHYTOCHROME B1* (*PHYB1*)-dependent manner. Importantly, genetic and molecular analyses combining a photoperiod-responsive tomato line carrying the wild *SP5G* allele from *S. pennellii,* together with *SlCOL1* and flowering-pathway mutants, revealed synergistic crosstalk among the photoperiodic *SlCOL1–SP5G* module, age-dependent pathway (mediated mainly by the microRNA156–*SlSBP* module), gibberellin (GA) pathway, and *SINGLE FLOWER TRUSS* (*SFT*) pathway. Mechanistically, we show that SP5G forms a complex with miR156-targeted SlSBP13 to directly regulate *SFT* expression, and that GA may interfere with SP5G activity. Together, these findings revealed a coordinated network that integrates multiple flowering signals to modulate both shared and pathway-specific targets. Our findings provide a significant advance in understanding the molecular regulation of tomato flowering and offer promising avenues for breeding strategies optimized for diverse environmental conditions and latitudes.

## INTRODUCTION

Photoperiod determines plant responses to the length of the day and is a crucial element in its physiology (Kinoshita and Richter, 2020). Day length is a reliable signal that depends on the Earth’s translational and rotational movements and is a robust cue for plants to choose the best time of the year to make important physiological and developmental decisions (Romero et al., 2024). It is so essential that traditionally plants have been classified according to photoperiod, namely long-day (LD) and short-day (SD) plants, as well as day-neutral plants, whose flowering is independent of the photoperiod (Garner and Allard, 1920; Andrés and Coupland, 2012). Despite being classified as day-neutral, cultivated tomato (*Solanum lycopersicum*) exhibits a residual photoperiodic response that delays flowering under LD conditions (Soyk et al., 2017). This photoperiodic response primarily depends on the activity of *SELF-PRUNING GENE 5* (*SP5G*) and *SP11B.1* (*FLOWERING LOCUS T-LIKE 1*, *FTL1*) genes, both members of the *CETS* (***CE****NTRORADIALIS*; ***T****ERMINAL FLOWER LOCUS 1*; and ***S****ELF-PRUNING*) gene family (Carmel-Goren et al., 2003; The Tomato Genome Consortium, 2012). Under LD, *SP5G* is highly expressed, resulting in delayed flowering due to the repression of *SINGLE FLOWER TRUSS* (*SFT*), another homologue of the mobile florigen *FLOWERING LOCUS T* (*FT*) (Soyk et al., 2017; Zhang et al., 2018). However, how SP5G represses *SFT* under LD remains unclear. In contrast, in SD, *SP11B.1* promotes *SFT* expression, inducing the floral transition (Song et al., 2020). Consequently, mutations in both *SP5G* and *SP11B.1* during domestication suppressed tomato photoperiodic flowering, enabling its cultivation across different latitudes (Song et al., 2020).

The expression of *SP5G* in LD was shown to be regulated by the photoreceptor *PHYTOCHROME B1* (*PHYB1*) (Cao et al., 2016), but how *SP5G* activity is modulated by PHYB1 is largely unknown. Several studies, primarily in *A. thaliana*, have unveiled the role of the BBX protein CONSTANS (CO) in regulating the photoperiod response. *Arabidopsis* CO forms a tripartite complex with PHYB and PHYTOCHROME-DEPENDENT LATE-FLOWERING (PHL), and it is the central hub of a complex photoperiodic regulatory network (Endo et al., 2013; Romero et al., 2024). Importantly, *Arabidopsis* CO is a direct transcriptional activator of *FT* (Tiwari et al., 2010; Robson et al., 2001; An et al., 2004). The tomato genome harbours 31 members of the *BBX* gene family (Lira et al., 2020). Among them, tomato *CONSTANS-like1* (*SlCOL1*), a *CO* homolog also referred to as *SlBBX3*, has been shown to regulate fruit yield and size by repressing *SFT* expression (Cui et al., 2022). In potato, flowering and tuberization are developmental processes that share key regulatory elements. Recently, the florigen *SELF PRUNING 3D* (*StSP3D*), an *SFT* homolog, was identified as a tuberization-inducing element in potato (Jing et al., 2023). Under LD conditions, tuberization is inhibited by *StCOL1* (a *SlCOL1* homolog), which directly binds to the *StSP5G* promoter and activates its expression. In turn, *StSP5G* suppresses *StSP6A* expression, an *FT* homolog primarily responsible for inducing tuber formation (Abelenda et al. 2016). Notably, both *StCOL1* and *StSP5G* exhibit peak expression in the morning in LD, a pattern also observed for *SP5G* and *SlCOL1* in tomato (Soyk et al., 2017; Yang et al., 2020; Zhang et al., 2018; Ben-Naim et al., 2006). However, the existence of an “external coincidence model” in tomato flowering (Valverde et al. 2004; Austen et al. 2017), and the precise role of *SlCOL1* in photoperiodic response, remain unclear.

*Arabidopsis* CO activity is regulated by distinct factors, including phytochromes (Valverde et al., 2004; Shim et al., 2017; Luo et al., 2022) and DELLA proteins (Xu et al., 2016; Wang et al., 2016). While *PHYA* and *PHYB* regulate CO protein stability (Valverde et al., 2004), DELLA, a key negative regulator of the gibberellin (GA) signalling, represses flowering in *Arabidopsis* under LD by inhibiting CO transcriptional activity (Hauvermale et al., 2012; Xu et al., 2016; Wang et al., 2016). Meanwhile, tomato DELLA (PROCERA or PRO) promotes flowering by interacting with the age-dependent pathway to activate *SFT* and other flowering-associated genes (Silva et al, 2019). The key components of the age-dependent pathway include the highly conserved microRNA156 (miR156) and its targets— members of the *SQUAMOSA PROMOTER BINDING PROTEIN–LIKE* (*SPL/SBP*) transcription factor family (Cardon et al., 1999; Wang et al., 2009; Rubio-Somoza and Weigel, 2011; Morea et al., 2016). In *Arabidopsis*, the miR156–*SPL/SBP* regulatory module functions downstream of *SUPPRESSOR OF OVEREXPRESSION OF CONSTANS 1* (*SOC1*) to initiate the floral transition. Interestingly, the *SOC1–SPL/SBP* regulatory module plays a key role in mediating GA signalling during the initiation of flowering under non-inductive SD conditions (Jung et al., 2012). On the other hand, under LD, miR156-targeted SPL3/4/5 interact with FLOWERING LOCUS D (FD), a basic leucine zipper transcription factor, and they directly bind to the promoters of *APETALA1*, *LEAFY*, and *FRUITFULL*, thus mediating their activation by the FT-FD flowering complex (Jung et al., 2016). Although the miR156-targeted *SlSBP13* promotes flowering in tomato by directly binding to the *SFT* promoter and activating its transcription (Cui et al., 2020), the interaction between the miR156–*SPL/SBP* module and the photoperiodic pathway in tomato remains unknown.

Understanding the crosstalk between flowering pathways is essential for advancing future research and agricultural practices (Han et al., 2021), but the interactions among the photoperiod-, GA-, and age-dependent flowering pathways in tomato and wild relatives remain poorly understood. Here, we employed a photoperiod-responsive tomato line harbouring a functional *SP5G* allele from a wild relative tomato species, and CRISPR mutants to demonstrate that SlCOL1 negatively regulates tomato flowering by directly activating *SP5G* expression in a *PHYB1*–dependent manner, thereby delaying flowering. Importantly, our genetic and molecular data indicated that *DELLA/PRO* and miR156-targeted *SlSBPs* act synergistically with *SP5G* and *SFT* to regulate flowering through both shared and pathway-specific target genes, particularly under LD conditions. Our findings reveal key aspects of the genetic regulation of tomato flowering, with potential applications for optimizing fruit production through fine-tuning of flowering time. Notably, precise regulation of flowering can alter plant growth habit, which in turn affects productivity and water use efficiency (WUE) (Vicente et al., 2015), traits that are particularly important under current climate change scenarios.

## RESULTS

### *SlCOL1* functions upstream of *SP5G* to control tomato photoperiodic flowering

Wild tomato relatives exhibit late flowering under LD conditions, a trait mainly regulated by *SP5G* (Soyk et al, 2017). However, how the *SP5G* activity is modulated in response to photoperiod remains uncertain. To investigate the regulatory components of photoperiodic flowering in cultivated tomato, we employed a photoperiod-responsive tomato line harbouring a functional *SP5G* allele from the photoperiodic-sensitive wild species *S. pennellii* (hereafter referred to as *SP5G^pen^*). The *SP5G^pen^* plants were generated by introgression of the *SP5G^pen^* allele into tomato cv. Micro-Tom (MT), using the introgression line (IL) 5-4 as a donor (Eshed and Zamir, 1995; Carmel-Goren et al., 2003). Genetic and molecular analyses confirmed the *SP5G^pen^* introgression into the MT background (Figure S1A; Siqueira et al, 2020), and that *SP5G^pen^* plants exhibited delayed flowering in LD conditions, producing more leaves to flowering and requiring more days to reach anthesis compared to MT (Figure S1B and 1C). These findings indicate that *SP5G* can affect both floral transition and floral bud development. As expected, LD conditions led to increased *SP5G* expression, especially in *SP5G^pen^* plants, which was accompanied by a decrease in *SFT* expression (Figure S1D and 1E), supporting the role of *SP5G* as an antiflorigen (Soyk et al., 2017).

While *CO* is a central regulator of *Arabidopsis* photoperiodic flowering, the potential interaction between *CO* and *SP5G* in tomato remains unclear. *CO* belongs to the BBX transcription factor family and promotes photoperiodic flowering by promoting *FT* expression (Robson et al., 2001; An et al., 2004; Tiwari et al., 2010; Valverde, 2011; Romero et al., 2024). In tomato, *SlCOL1*, a *CO* homolog, was recently shown to act as a repressor of fruit yield by downregulating *SFT* (Cui et al, 2022) and to contribute to inflorescence adaptation under high temperature (Sun et al., 2024). However, its role as a regulatory component of tomato photoperiodic flowering is still unexplored. To evaluate this possibility, we assessed *SlCOL1* expression in leaves of *SP5G^pen^* and MT plants growing under SD and LD conditions. Regardless of the photoperiod, we did not observe differences in *SlCOL1* transcript levels between the photoperiod-sensitive (*SP5G^pen^*) and -insensitive (MT) plants. However, we detected a significant increase in *SlCOL1* transcript level under LD in both genotypes (Figure 1A), suggesting that *SlCOL1* may act upstream of *SP5G* in the photoperiodic flowering pathway.

**Figure 1.**
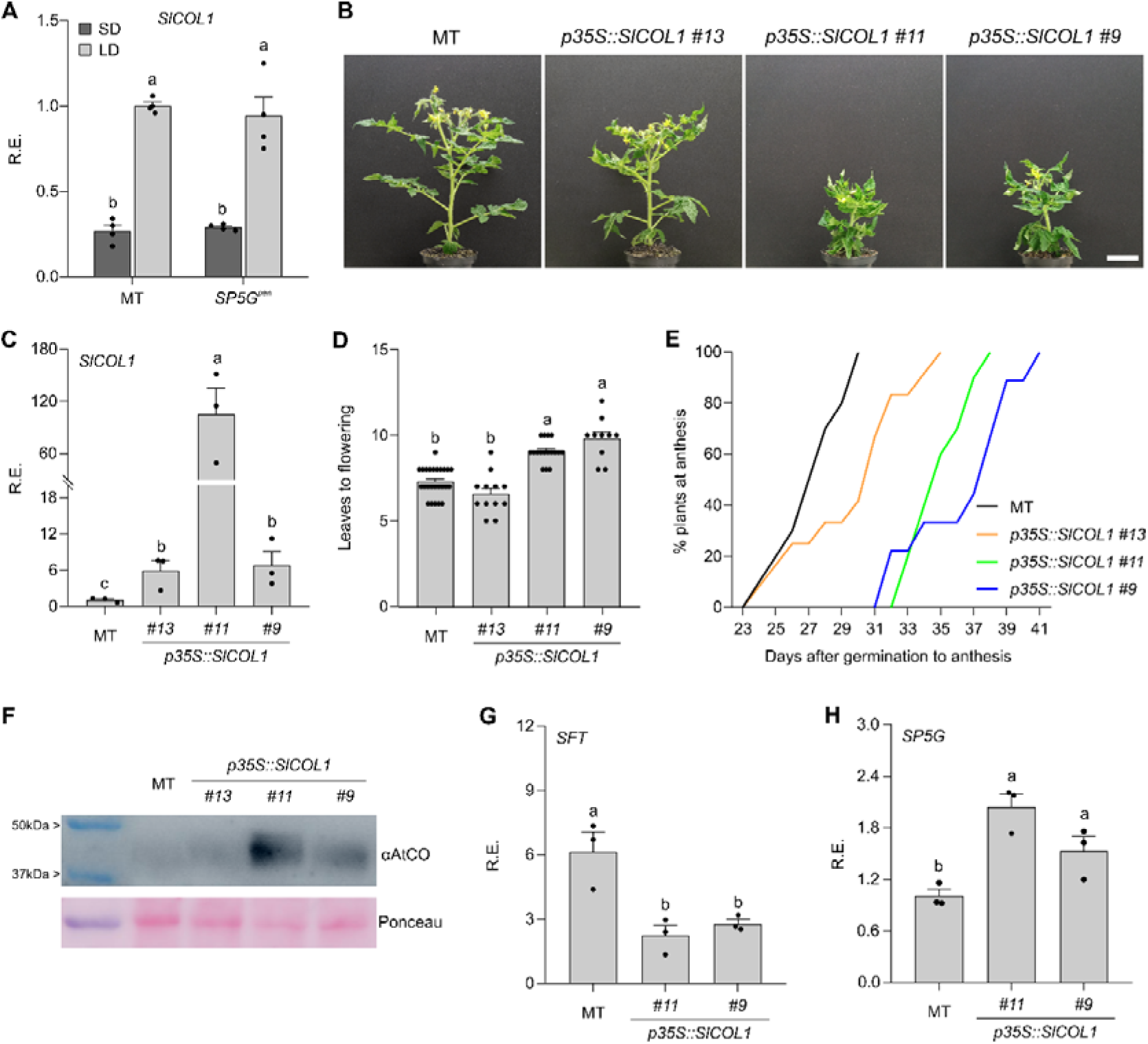
*SlCOL1* overexpression represses tomato flowering. **(A)** Relative expression (R.E.) of *SlCOL1* in photoperiod-insensitive (MT) and -sensitive (*SP5G^pen^*) plants under short day (SD, 10/14h, dark grey bars) and long day (LD, 14/10h, light grey bars) conditions (ZT = 4; mean ± SEM; Tukey’s test, *p<0.05*; n = 3 samples). **(B)** Representative images of 40-day-old transgenic plants overexpressing *SlCOL1*. Bar = 5cm. **(C)** Relative expression of *SlCOL1* in three selected independent lines (ZT = 4; *p35S::SlCOL1 #9*, *#11*, and *#13*). **(D)** Leaves to flowering of MT and *p35S::SlCOL1* lines in LD (mean ± SEM; Tukey’s test, *p<0.001*; n = 10-25). **(E)** Percentage of plants at anthesis after germination in LD conditions for MT (black line) and *p35S::SlCOL1* plants (n = 10-25). Blue, green, and orange lines represent *p35S::SlCOL1* plants from lines *#9*, *#11*, and *#13*, respectively. (**F)** Immunoblot analysis of SlCOL1 protein accumulation in MT and *p35S::SlCOL1* seedlings (10 DPG, ZT = 4). The membrane was incubated with αAtCO (anti-AtCO). Ponceau staining was used as a loading control. The ladder is shown on the left. (**G and H)** Relative expression of *SFT* (**G**) and *SP5G* (**H**) in MT and *p35S::SlCOL1* lines #9 and #11 in LD (ZT = 4; mean ± SEM; Distinct letters indicate significant differences according to Tukey’s test, *p<0.05*; n = 3 samples). Each dot represents individual data.

To test this hypothesis, we generated *Slcol1* mutants in the MT background using CRISPR/Cas9 technology (referred to as *Slcol1^CR^*) to edit out the middle domain between the BBX and CCT domains (Valverde, 2011). We obtained two independent *slcol1^CR^*mutant alleles in the MT background (namely #1 and #2), and neither of them exhibited significant changes in flowering (Figure S2A-S2C. This lack of strong flowering phenotypes in *slcol1^CR^* mutants was previously reported and suggested redundancy with other closed-related tomato *BBXs* such as *SlCOL2* and *SlCOL3* (Lira et al., 2020; Cui et al., 2022). To further test this conjecture, we introduced the *SP5G^pen^*into *slcol1^CR^*#1 plants. The flowering time of the resulting *SP5G^pen^ slcol1^CR^*#1 plants closely resembled that of *SP5G^pen^* (Figure S2A-S2C), indicating partial functional redundancy among *COL* genes in cultivated tomato.

Given that cultivated tomato has undergone a long process of domestication, in addition to functional redundancy among *COL* genes, other components of the photoperiodic pathway may have been altered (Song et al., 2020). Because wild relatives exhibit stronger photoperiodic responses (Soyk et al., 2017), we similarly generated *col1* mutants in *S. pimpinellifolium*, which is the closest wild relative of cultivated tomato and harbours a photoperiod-responsive *SP5G* allele (Aflitos et al., 2014; Soyk et al., 2017). We generated three distinct CRISPR mutants (referred here to as *Spcol1^CR^*) and evaluated their flowering time and days to anthesis. Mutations in the middle domain of the *SpCOL1* gene accelerated flowering in LD conditions (Figure S2D-S2F), indicating that *SpCOL1* plays a crucial role in repressing photoperiodic flowering in wild species. Importantly, *SpSP5G* transcript levels were reduced in *Spcol1^CR^* seedlings, suggesting that SpCOL1 is required for *SP5G* activation in *S. pimpinellifolium* under LD (Figure S3A).

To further investigate the role of *SlCOL1* in photoperiodic responses in tomato, we generated transgenic MT plants overexpressing the *SlCOL1* coding sequence (CDS), and selected three independent lines (*p35S::SlCOL1 #9, #11*, and *#13*) for further analyses (Figure 1B). Although all evaluated lines showed high *SlCOL1* expression (Figure 1C), lines #9 and #11 exhibited modifications in vegetative architecture, such as leaf-curling and reduced stature. Importantly, *SlCOL1*-overexpression lines #9 and #11 exhibited an increased number of leaves until the first inflorescence, and required more days to anthesis (Figure 1B; 1D-1E). These findings suggested that high levels of *SlCOL1* impact both vegetative and reproductive growth in tomato. Interestingly, line #13 exhibited phenotypes comparable to the MT plants (Figure 1B), which might indicate modifications at the protein level rather than at the transcriptional levels. To explore this possibility, we evaluated SlCOL1 protein levels in 10-day post-germination (10 DPG) seedlings. *In silico* predictions indicate that *SlCOL1* encodes a protein of 391 aa (∼ 43.4 kDa) (https://www.bioinformatics.org/sms/prot_mw.html). To detect SlCOL1 protein in Western blot assays, we employed an *Arabidopsis* CO antibody (αAtCO), which was generated against the middle domain of the CO protein (Valverde et al., 2004). Our results showed that the αAtCO was able to detect tomato COL1 protein in MT and *S. pimpinellifolium* (Figure S3B-S3C). As expected, *Slcol^CR^* and *Spcol1^CR^* mutant seedlings exhibited lower COL1 protein levels when compared with the respective controls (Figure S3B-S3C). Nevertheless, we cannot rule out the possibility that other CO-like (COL) proteins may also be recognized by this antibody. Importantly, *p35S::SlCOL1* lines #9 and #11 accumulated more SlCOL1 protein than MT and line #13 (Figure 1F), likely accounting for the developmental differences between line #13 and the other overexpressing lines.

A previous study demonstrated that *SlCOL1*-overexpressing plants exhibit reduced *SFT* expression (Cui et al., 2022), which was also observed in our plants (Figure 1G). The increased expression of *SlCOL1* in LD (Figure 1A) suggested that it might be important to modulate tomato photoperiodic responses. Potato (*S. tuberosum*) *COL1*, a *SlCOL1* homolog, inhibits tuberization specifically in LD conditions by directly activating *StSP5G* expression (Abelenda et al. 2016). Given the high level of synteny between the tomato and potato genomes (The Tomato Genome Consortium, 2012), we hypothesized that *SlCOL1* represses tomato flowering in LD by directly regulating *SP5G*. To test this hypothesis, we first evaluated *SP5G* expression in *p35S::SlCOL1* lines #9 and #11 grown in LD. The results revealed a significant increase in *SP5G* transcript levels in both overexpressing lines (Figure 1H), suggesting that SlCOL1 activates *SP5G* in tomato under LD. Given that *p35S::SlCOL1* lines #9 and #11 displayed similar phenotypes, and that line #11 could not be stabilized at a homozygous state, we selected line #9 (hereafter referred to only as *p35S::SlCOL1*) for further analyses.

### SlCOL1 binds to the *SP5G* promoter to activate its expression

To further elucidate the role of *SlCOL1* in photoperiodic flowering, we introduced the *SP5G^pen^* allele into *p35S::SlCOL1* plants, generating the *p35S::SlCOL1 SP5G^pen^* genotype. In LD, *p35S::SlCOL1 SP5G^pen^* plants showed vegetative phenotypes more similar to *p35S::SlCOL1* plants (Figure 2A). By contrast, *p35S::SlCOL1 SP5G^pen^* plants displayed vegetative phenotypes similar to both parental lines under SD conditions (Figure S4A). These observations suggested that under LD conditions, elevated levels of *SlCOL1* and *SP5G^pen^* modulate independent pathways during vegetative development. However, we observed an additive effect on flowering time in LD, with an additional increase in the number of leaves to the first inflorescence and in the days to anthesis in *p35S::SlCOL1 SP5G^pen^* when compared to their parental lines (Figure 2A-2C). Notably, *SlCOL1-*overexpressing plants also exhibited a higher number of leaves and a longer time to anthesis in SD, particularly in those plants carrying the *SP5G^pen^*allele (Figure S4). Our results indicate that, in addition to repressing *SFT* (Cui et al., 2022), high levels of SlCOL1 protein delayed tomato flowering in an *SP5G*-dependent manner under both LD and SD conditions (Figure 2A-2C; Figure S4). Importantly, introducing the *SP5G^pen^* allele into *p35S::SlCOL1* lines #11 and #13 altered vegetative and flowering traits only in line #11, consistent with the distinct protein levels observed in these transgenic lines (Figure S5).

**Figure 2.**
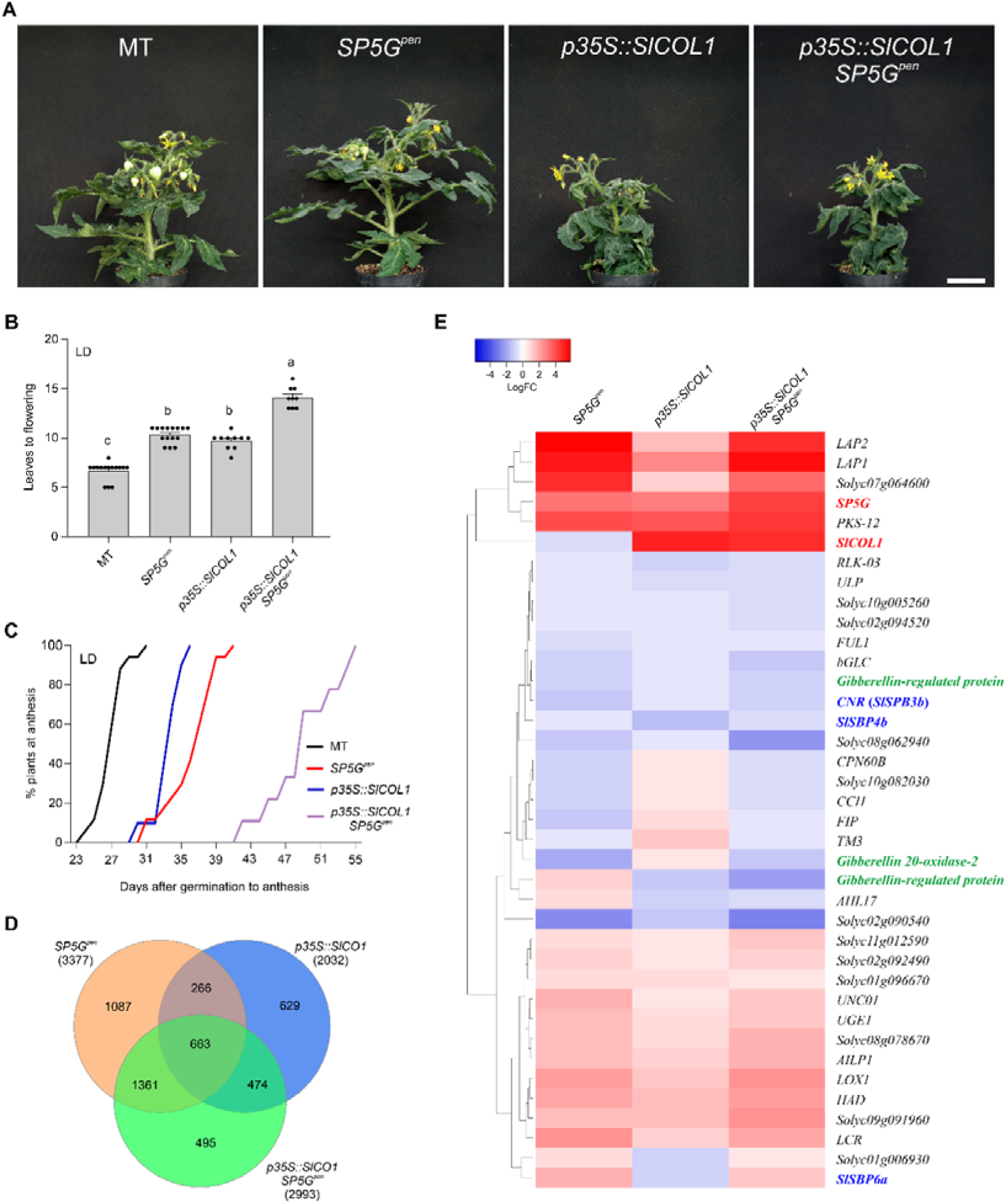
High levels of *SlCOL1* and *SP5G* repress tomato flowering in LD. **(A)** Representative images of 60-day-old plants of MT, *SP5G^pen^*, *p35S::SlCOL1*, and *p35S::SlCOL1 SP5G^pen^* cultivated in LD (14h/10h). Bar = 5cm. **(B)** Leaves to flowering of MT, *SP5G^pen^*, *p35S::SlCOL1*, and *p35S::SlCOL1 SP5G^pen^* in LD. Data are expressed as mean ± SEM (Tukey’s test, *p<0.001*; n = 9-15). Each dot represents individual data. **(C)** Percentage of plants at anthesis after germination in LD conditions for MT (black line), *SP5G^pen^* (red line), *p35S::SlCOL1* (blue line), and *p35S::SlCOL1 SP5G^pen^* (purple line) (n = 8-15). **(D)** Venn diagram showing the number of differentially expressed genes (DEGs) shared among *SP5G^pen^*, *p35S::SlCOL1*, and *p35S::SlCOL1 SP5G^pen^* two-week-old seedlings cultivated in LD. Numbers below the genotypes indicate the total number of DEGs identified in each genotype compared to MT. **(E)** Expression profiles of 38 shared DEGs among *SP5G^pen^*, *p35S::SlCOL1*, and *p35S::SlCOL1 SP5G^pen^* seedlings compared to MT. Genes are associated with photoperiod (red), age-related (blue), and gibberellin (green) flowering pathways, as well as known floral transition markers (black, Meir et al., 2021). Scale bar represents Log_2_ fold change values.

To better understand the molecular mechanisms by which *SlCOL1* and *SP5G^pen^* modulate tomato flowering in non-inductive conditions (LD), we used RNA sequencing (RNA-seq) to monitor changes in gene expression in two-week-old MT, *SP5G^pen^*, *p35S::SlCOL1*, and *p35S::SlCOL1 SP5G^pen^* seedlings grown in LD. The RNA-seq analysis identified 3377, 2032, and 2993 differentially expressed genes (DEGs) in *SP5G^pen^*, *p35S::SlCOL1*, and *p35S::SlCOL1 SP5G^pen^*, respectively (Figure 2D; Table S1-S3). The difference in DEG number between *SP5G^pen^* and *p35S::SlCOL1* seedlings might account for their distinct vegetative phenotypes (Figure 2A). Importantly, we found 663 DEGs shared among the three genotypes (Figure 2D; Table S4), including genes associated with the age-dependent and GA pathways, as well as known flowering markers (Figure 2E; Table S4; Meir et al., 2021). As expected, *SP5G* was one of the most upregulated genes in *SP5G^pen^*, *p35S::SlCOL1*, and *p35S::SlCOL1 SP5G^pen^* seedlings. *SP5G* expression was ∼8-fold higher in *SP5G^pen^* and *p35S::SlCOL1* plants compared with MT, whereas *p35S::SlCOL1 SP5G^pen^* showed nearly a 20-fold increase (Figure 3A). We also quantified *SP5G* transcript accumulation in MT, *SP5G^pen^*, *p35S::SlCOL1*, and *p35S::SlCOL1 SP5G^pen^* seedlings in SD by qRT-PCR. Consistent with their later flowering phenotypes in SD conditions (Figure 1D-1F), *p35S::SlCOL1* and *p35S::SlCOL1 SP5G^pen^* plants showed ∼33-and ∼40-fold increase in *SP5G* expression compared to MT, respectively (Figure 3B), confirming that higher *SlCOL1* levels promote *SP5G* transcription and repress flowering under inductive and non-inductive conditions.

**Figure 3.**
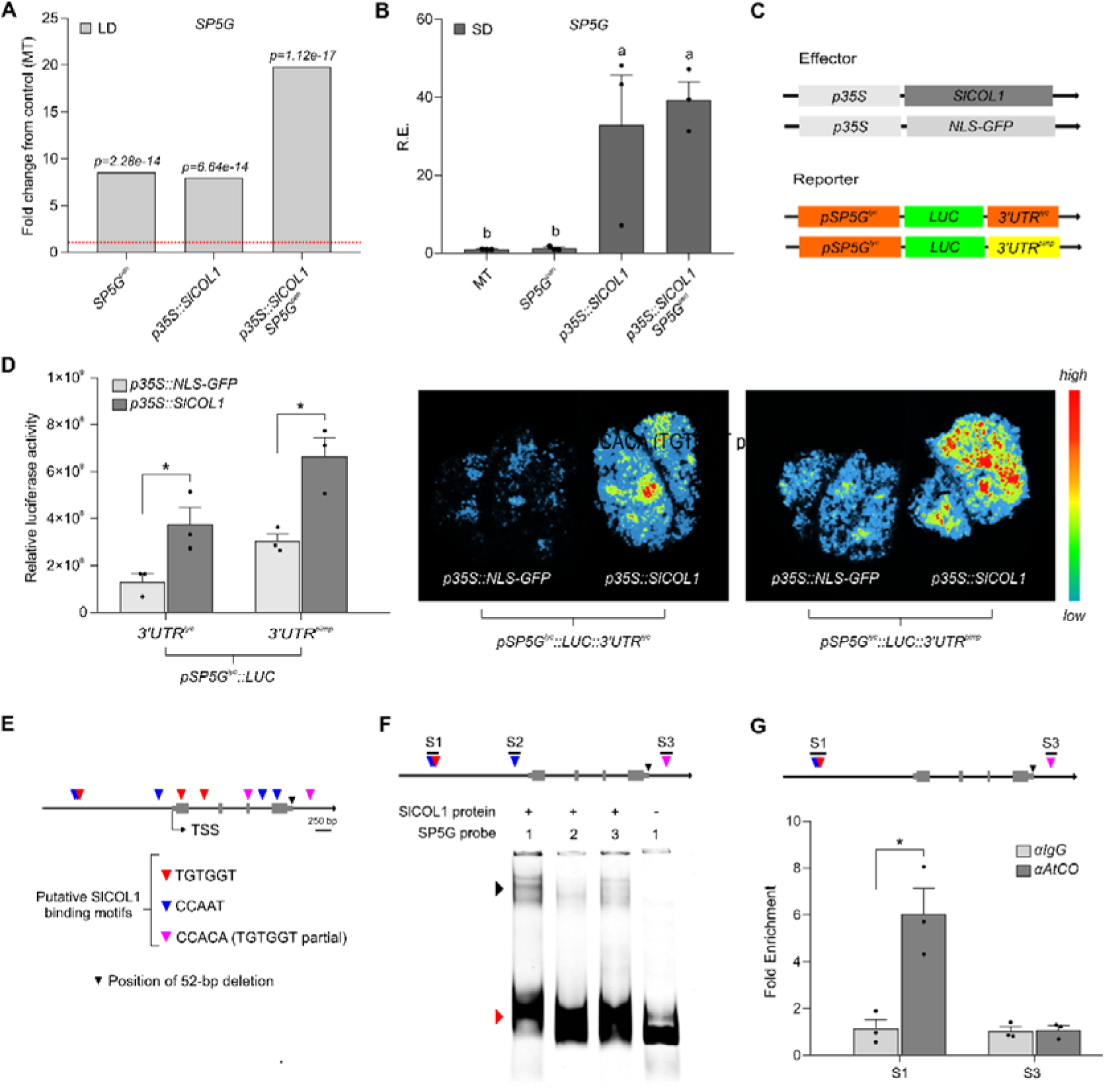
SlCOL1 binds to the *SP5G* promoter and activates its expression. **(A)** RNA-seq data (Fold-change) of *SP5G* in *SP5G^pen^*, *p35S::SlCOL1*, and *p35S::SlCOL1 SP5G^pen^* two-week-old seedlings in LD (14/10h) conditions. *p-*values are shown above the bars. **(B)** Relative expression (R.E.) of *SP5G* in MT, *SP5G^pen^*, *p35S::SlCOL1*, and *p35S::SlCOL1 SP5G^pen^* seedlings in SD (10/14h) (ZT = 4; mean ± SEM; n = 3). Distinct letters indicate significant differences according to Tukey’s test (*p<0.05*). **(C)** Schematic representation of effector and reporter constructs used in the transient expression assay. Reporter constructs harbouring the tomato *SP5G* promoter (*pSP5G^lyc^*) drive the luciferase (*LUC*) expression fused to either the tomato (*3’UTR^lyc^*) or *S. pimpinellifolium* (*3’UTR^pimp^*) 3’UTRs. **(D)** Transient activation of *pSP5G^lyc^::LUC::3’UTR^lyc^* and *pSP5G^lyc^::LUC::3’UTR^pimp^* reporter constructs in *N. benthamiana* leaves. Three leaves were co-infiltrated with the reporter constructs plus either *p35S::SlCOL1* or *p35S::NLS-GFP* effector constructs. Right: quantification of relative luciferase activity using NEWTON 7.0 imaging software (https://animalab.eu/newton-7-0-in-vivo-imaging-system). Values are mean ±SEM (Student’s *t*-test, *p<0.05*, n = 3). Left: Representative bioluminescence images captured using the NEWTON 7.0 CCD imaging system. Scale represents the luciferase activity intensity. **(E)** Schematic representation of the *SP5G* locus showing putative SlCOL1-binding motifs. Grey boxes represent exons. TSS indicates the position of the transcription start site in the locus. Colored triangles indicate putative SlCOL1 binding motifs, including TGTGGT (red), CCAAT (blue), and CCACA (magenta; partial TGTGGT motif). The black triangle marks the position of the 52-bp deletion. **(F)** EMSA to test the ability of SlCOL1 to interact with distinct regulatory regions of the *SP5G* gene. Black arrowhead indicates the protein-probe complex, whereas the red arrowhead indicates the free probe. **(G)** ChIP fold enrichment of DNA fragments harboring *SlCOL1* binding motifs immunoprecipitated with αAtCO and αΙgG (IgG antibody; control) from *p35S::SlCOL1* seedlings and quantified by qPCR (mean ± SEM; Student’s *t*-test, p<0.05, n=3). The schematic representations at the top of panels (**F**) and (**G**) indicate the relative positions of the tested probes and fragments at the *SP5G* locus, respectively. Each dot represents individual data.

The lower expression of the *SP5G* allele under LD in cultivated tomato compared with wild relatives is caused by a 52-bp deletion in its 3’UTR, which acts as a transcriptional enhancer (Zhang et al., 2018). Thus, to better understand the role of *SlCOL1* in the transcriptional regulation of *SP5G*, we generated two reporter constructs in which the tomato *SP5G* promoter (*pSP5G^lyc^*) drove the luciferase (*LUC*) expression: one carrying the tomato 3’UTR (*pSP5G^lyc^*::*LUC::3’UTR^lyc^*) and the other carrying the *S. pimpinellifolium* 3’UTR (*pSP5G^lyc^*::*LUC::3’UTR^pimp^*) (Figure 3C). We then co-expressed these constructs with the effector constructs, *p35S::SlCOL1* or *p35S::NLS-GFP*, in *Nicotiana benthamiana* leaves. Regardless of the reporter construct, LUC activity was significantly increased in the presence of SlCOL1, with higher activity observed when the 3′ UTR from *S. pimpinellifolium* was present (Figure 3D), as expected, since it contains the 52-bp region that acts as an enhancer (Zhang et al., 2018). Collectively, these findings provide evidence that *SlCOL1* delays tomato flowering through a direct transcriptional activation of *SP5G*.

In potato, StCOL1 binds to a conserved TGTGGT motif in the *StSP5G* promoter, inducing its expression (Abelenda et al. 2016). Similarly, in *Arabidopsis*, AtCO regulates *FT* expression by binding to the CO-responsive CORE (CCACA, a TGTGGT partial motif) and CCAAT-box motifs (Wenkel et al., 2006; Tiwari et al., 2010; Gnesutta et al., 2017). By screening the *SP5G* locus, we identified 10 putative *SlCOL1*-binding motifs: three TGTGGT, five CCAAT, and two CCACA. Among those, four motifs are in the promoter (with three clustered approximately 1.4 kb upstream of the start codon), five within the open reading frame (ORF), and one in the 3’ UTR region (Figure 3E). Based on the data of our LUC assays (Figure 3D) and the region of the *SlCOL1*-binding motifs (Figure 3E), we selected two regions in the *SP5G* promoter (S1 and S2) and one in the 3’UTR (S3, which not included the 52-bp deletion) to generate probes for Electrophoretic Mobility Shift Assays (EMSA) using *in-vitro* produced SlCOL1 protein. Our results revealed a slight shift in the mobility of the S1 and S3 probes when incubated with SlCOL1, suggesting that the SlCOL1 affinity for these regions may require a cofactor or additional regulatory elements to enhance binding stability (Figure 3F). Remarkably, the EMSA data revealed similar SlCOL1 binding to the *SP5G* promoter (S1 probe) as previously observed for the *SFT* promoter region (S1 probe; Figure S6; Cui et al., 2022), reinforcing the affinity of SlCOL1 for the *SP5G* promoter. To further validate the binding of SlCOL1 to the *SP5G* promoter, we performed Chromatin Immunoprecipitation (ChIP)–qPCR assay for regions S1 and S3 in *p35S::SlCOL1* plants, using the αAtCO antibody and seedlings at 10 DPG as samples. Seedlings were grown in LD conditions, and samples were collected at ZT4, when *SP5G* expression is higher (Zhang et al., 2018). The immunoprecipitated fraction with αAtCO was significantly enriched only in the S1 region when compared with the control (Figure 3G), indicating that SlCOL1 regulates SP5G mainly through its promoter region, similarly to what was shown by Song et al. (2025). The lack of enrichment in the S3 region may be due to the absence of relevant cis-regulatory elements required for SlCOL1 association in this region. Collectively, our data provide robust evidence that SlCOL1 negatively regulates tomato flowering by binding to the *SP5G* promoter, thereby facilitating its expression under long-day conditions.

### *phyB1* mutation is epistatic to *SP5G^pen^* and *p35S::SlCOL1* late flowering phenotypes

The tomato *phyB1* mutant exhibits reduced expression of *SP5G* under LD conditions (Figure 4A; Cao et al., 2016), indicating that this phytochrome modulates *SP5G* expression. To investigate how *PHYB1* modulates *SP5G* activity at the genetic level, we introduced the *phyB1* allele into *SP5G^pen^* plants and evaluated *phyB1 SP5G^pen^* flowering time. Our results showed that the *phyB1* mutation rescued the late flowering phenotype of *SP5G^pen^* plants in LD (Figure S7A-S7C), indicating that, independent of the *SP5G* allele (cultivated or wild), the *phyB1* mutation overrode the *SP5G* effect on flowering time. This suggests that the *phyB1* mutation is epistatic to the *SP5G^pen^* allele. Interestingly, under SD conditions, *phyB1* plants flowered later than MT (Figure S7D-S7F), indicating that *PHYB1* functions as a repressor of flowering under LD, but as a promoter under SD conditions.

**Figure 4.**
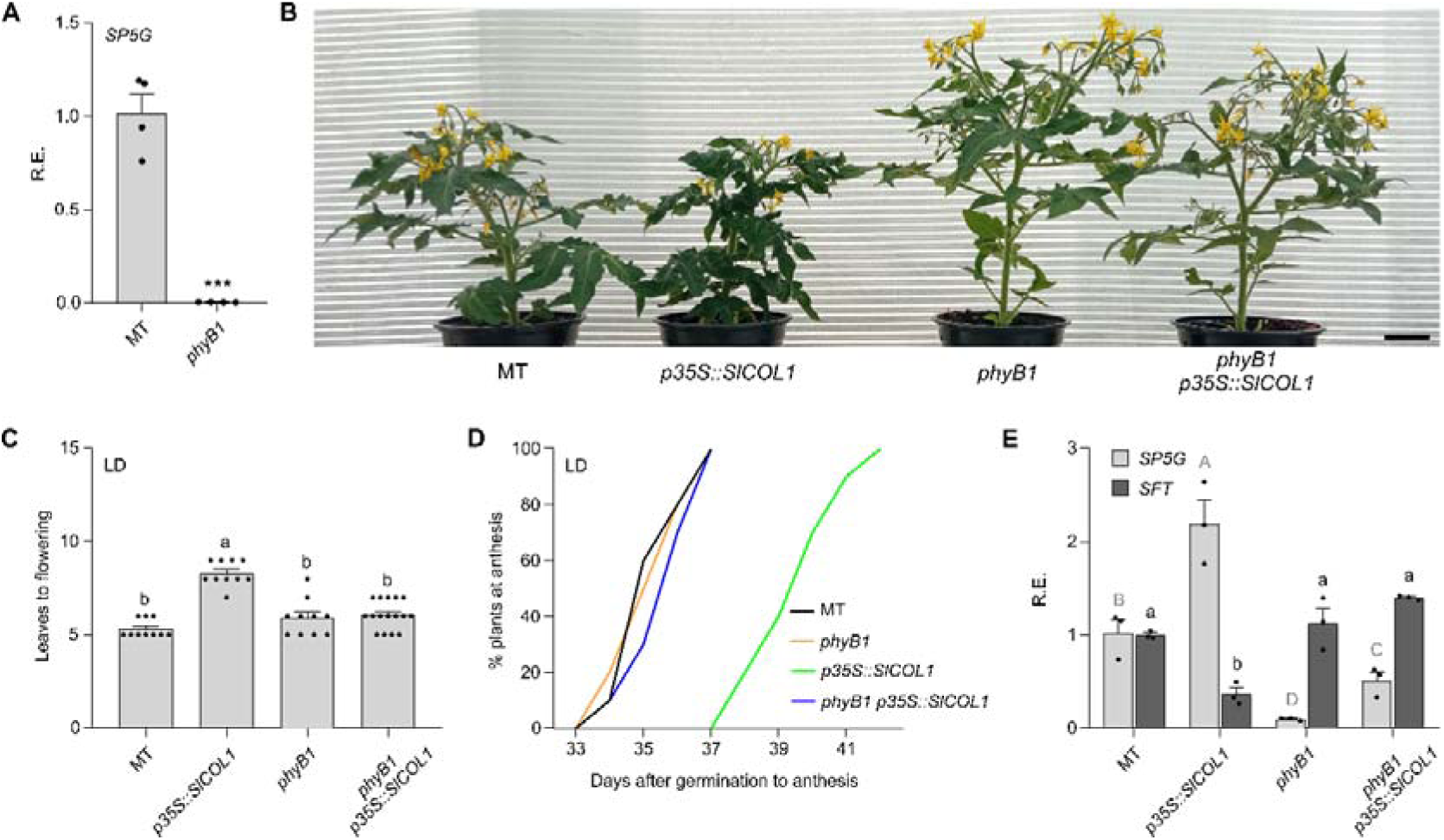
The *phyB1* mutation rescues the late flowering phenotype of *p35S::SlCOL1* plants. **(A)** Relative expression (R.E.) of *SP5G* in MT and *phyB1* seedlings (at 10 days post-germination or DPG) under LD conditions (ZT = 4; mean ± SEM; Student’s *t*-test, *p<0.01*; n = 4). **(B)** Representative images of 45-day-old MT, *phyB1*, *p35S::SlCOL1*, and *phyB1 p35S::SlCOL1* plants cultivated in LD. Bar = 5cm. **(C and D)** Leaves to flowering (**C**) and percentage of plants at anthesis after germination in LD conditions (**D**) for MT (black line), *phyB1* (orange line), *p35S::SlCOL1* (green line), and *phyB1 p35S::SlCOL1* (blue line) plants (n = 10-16). **(E)** Relative expression (R.E.) of *SP5G* and *SFT* in 10-day-old MT, *phyB1*, *p35S::SlCOL1*, and *phyB1 p35S::SlCOL1* plants cultivated in LD (ZT = 4; mean ± SEM; Tukey’s test, *p<0.05*; n = 3). Different uppercase and lowercase letters above the bars indicate significant differences in *SP5G* and *SFT* expression among genotypes, respectively. Each dot represents individual data.

Unlike *Arabidopsis phyB* mutant, which exhibits increased abundance of CO protein in the morning (Valverde et al., 2004), transgenic potato *phyB*-RNAi lines show reduced StCOL1 abundance under LD and SD conditions. These findings suggest that StPHYB is essential for the post-transcriptional regulation of *StCOL1* (Abelenda et al., 2016). Considering these findings and our data showing the positive regulatory role of PHYB1 and SlCOL1 on *SP5G* activity (Figure 2 and 3; Figure S7), we hypothesized that tomato PHYB1 also positively regulates SlCOL1 activity. To test this possibility at the genetic level, we introduced the *phyB1* allele into the *p35S::SlCOL1* background, generating the *phyB1 p35S::SlCOL1* plants. *phyB1 p35S::SlCOL1* plants displayed flowering time similar to that of MT and *phyB1*, effectively rescuing the late flowering phenotype observed in *p35S::SlCOL1* plants (Figure 4B-4D).

Additionally, *phyB1 p35S::SlCOL1* plants exhibited a vegetative architecture similar to that of *phyB1* (Figure 4B). These findings indicated that the *phyB1* mutation is also epistatic to the *SlCOL1* overexpression. To further explore the interaction between these genes, we investigated whether *PHYB1* is required for *SlCOL1* activity in tomato, as previously reported in potato (Abelenda et al., 2016). To this end, we quantified the transcript levels of *SlCOL1*-targeted *SP5G* and *SFT* in *phyB1 p35S::SlCOL1* plants. Consistent with their flowering phenotype (Figure 4B-4D), the *phyB1* mutation attenuated *SP5G* expression while enhancing *SFT* expression in the *p35S::SlCOL1* background (Figure 4E). Our results suggest that PHYB1 may modulate SlCOL1 activity through post-transcriptional mechanisms, although this possibility requires further investigation.

### Age- and photoperiodic-dependent pathways synergistically interact to orchestrate tomato flowering

We previously reported that miR156-overexpressing plants (*miR156OE*) and the *procera/della* (*pro*) mutant exhibit delayed flowering. Importantly, genetic and molecular data indicated synergistic interactions between the age-dependent miRNA156-*SlSBP* module and GA pathways in tomato flowering (Silva et al., 2019). Our RNA-seq analysis of two-week-old seedlings grown under LD conditions identified several differentially expressed miR156-targeted *SlSBP* genes, including *CNR*, *SlSBP3a*, *SlSBP4b*, *SlSBP6a*, *SlSBP6c*, and *SlSBP10* (Figure 2E; Table S1-S4). These *SlSBPs* represent major SPL/SBP clades in tomato (Salinas et al., 2012), including *SlSBP3a*, which has been previously associated with the regulation of floral transition (Silva et al., 2019; Meir et al., 2021). *SlSBP4b* was not previously reported by Salinas et al. (2012), but it contains a conserved miR156 recognition site in its 3′UTR and clusters within the flowering-associated SPL4 clade (Figure S8; Jung et al., 2016). These findings suggested a crosstalk between the photoperiod and age pathways in tomato, with the miR156–*SlSBP* module acting downstream of the photoperiodic pathway. This notion is further supported by the near wild-type transcript levels of *SlCOL1* and *SP5G* observed in miR156-overexpressing seedlings (Figure S9A-S9B).

Given that *SP5G* functions downstream of *SlCOL1* and *PHYB1* in tomato (Figure 2-3; Figure S7), we explored the potential genetic interaction between *SP5G* and the miR156-*SlSBP* module. To that end, we introduced the *SP5G^pen^* allele into *miR156OE* plants. Under LD, we observed a synergistic effect on flowering time in the double *miR156OE SP5G^pen^* compared with parental lines. While *miR156OE* and *SP5G^pen^* produced ∼13 and ∼10 leaves to flowering, respectively, the *miR156OE SP5G^pen^* exhibited more than 18 leaves. Additionally, the time to anthesis was significantly extended in *miR156OE SP5G^pen^* compared with the other genotypes (Figure 5A-5C). In contrast, under SD conditions, *miR156OE SP5G^pen^* plants flowered similarly to *miR156OE*. Likewise, *SP5G^pen^* displayed a flowering pattern comparable to MT (Figure S10). Together, these findings indicated that under LD conditions, where both SP5G and miR156 repress flowering, the PHYB1–SlCOL1–SP5G and miR156–*SlSBP* regulatory modules act synergistically. Conversely, under SD conditions, the miR156– *SlSBP* module assumes a dominant role in controlling tomato flowering.

**Figure 5.**
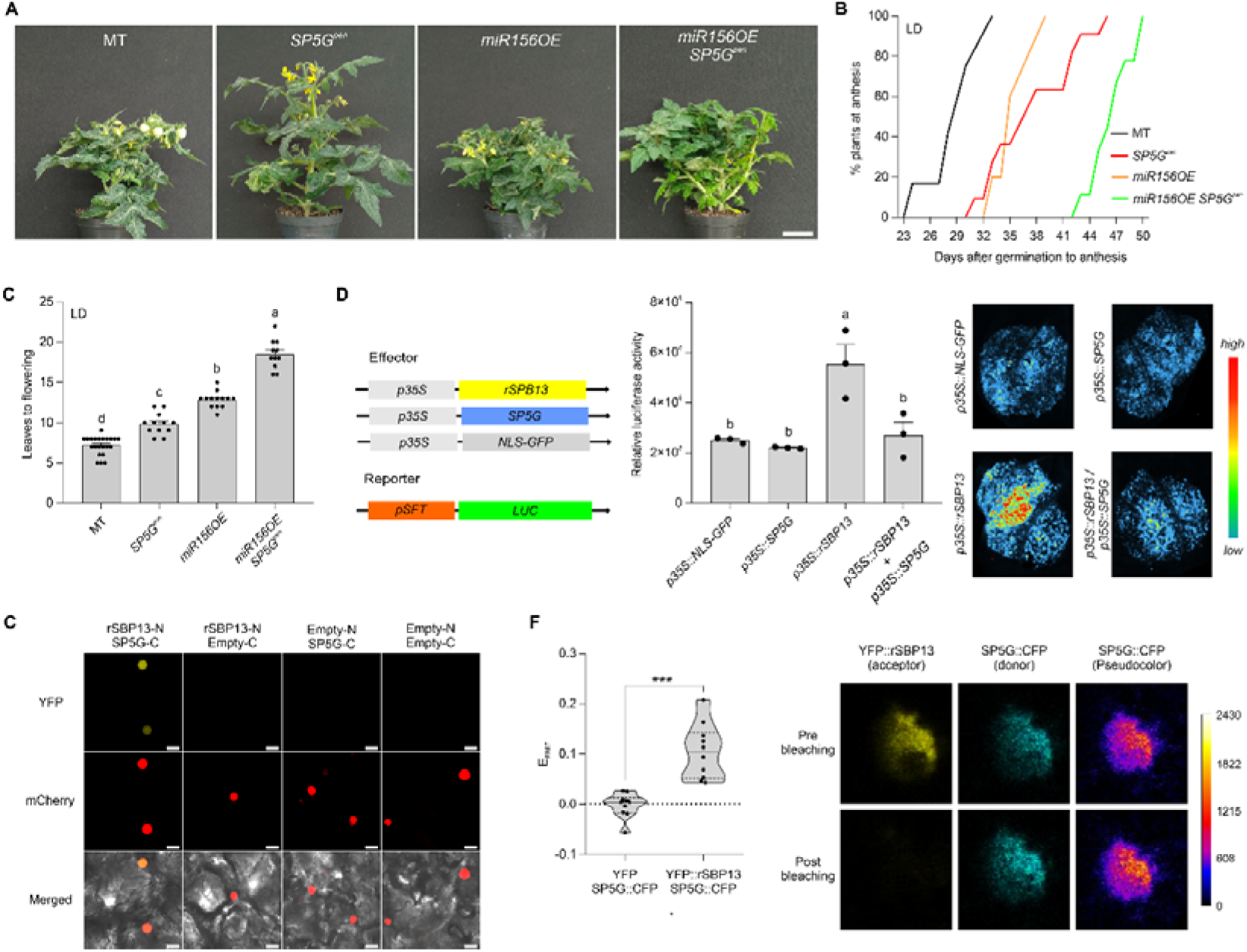
SP5G attenuates *SFT* activation by SlSBP13. **(A)** Representative pictures of 60-day-old plants of MT, *SP5G^pen^*, *miR156OE*, and *miR156OE SP5G^pen^* cultivated in LD (14h/10h). Bar = 5cm. **(B and C)** Percentage of plants at anthesis after germination in LD conditions (**B**) and leaves to flowering (**C**) for MT, *SP5G^pen^*, *miR156OE*, and *miR156OE SP5G^pen^* plants. In (**B**), black, red, orange, and green lines represent MT, *SP5G^pen^*, *miR156OE*, and *miR156OE SP5G^pen^* genotypes, respectively. The data are expressed as mean ± SEM (n = 11-22). Distinct letters indicate significant differences according to Tukey’s test (*p<0.001*). **(D)** Transient activation of *pSFT::LUC* reporter constructs in *N. benthamiana* leaves. Three leaves were co-infiltrated with *p35S::rSBP13*, *p35S::SP5G*, and/or *p35S::NLS-GFP* effector constructs. *rSBP13* represents a miR156-resistant version of *SlSBP13*. Left: schematic representation of effector and reporter constructs used in this assay. Middle: quantification of relative luciferase activity using NEWTON 7.0 imaging software. Values are mean ±SEM (n = 3). Distinct letters indicate significant differences according to Tukey’s test (*p<0.05*). Each dot represents an individual data. Right: representative bioluminescence images captured using the NEWTON 7.0 CCD imaging system. **(E)** Bimolecular fluorescence complementation (BiFC) assay using *N. benthamiana* leaves infiltrated with agrobacteria strain GV3101. N and C represent the N-terminal and C-terminal fragments of the YFP protein, respectively. Empty indicates an empty vector. Combinations of rSBP13-N with Empty-C, Empty-N with SP5G-C, and Empty-N with Empty-C were used as negative controls. AtWWP1::mCherry was used as a nuclear marker. Bright field and merged images are also shown. Scale bars = 20 μm. **(F)** Förster Resonance Energy Transfer (FRET) acceptor photobleaching measurements between SP5G and rSBP13. FRET efficiency (E_FRET_) was calculated using the equation: E_FRET_ = 1-F_DA_/F_D_, where F_DA_ and F_D_ indicate CFP signal before and after photobleaching, respectively. YFP alone co-infiltrated with SP5G::CFP was used as the negative control. Values are means ±SEM (Mann–Whitney U test, *p<0.001*, n = 10 cells). Representative images of YFP::rSBP13 and SP5G::CFP signals before and after photobleaching are shown (right). Images of the negative control are presented in Figure S11. To improve visualization, CFP fluorescence before and after photobleaching is shown using pseudocolour. A colour scale indicates the pixel intensity values. Each dot represents individual data.

High levels of *SP5G* lead to reduced *SFT* expression (Soyk et al., 2017, Figure S1E), whereas miR156-targeted *SlSBP13* promotes tomato flowering by activating *SFT* transcription (Cui et al., 2020). *SlSBP13* was not among the DEGs in our transcriptomic data (Table S1–S4), suggesting that it is not regulated by the SlCOL1–SP5G module at the transcript level. Thus, we hypothesized that SP5G may act as a transcriptional cofactor and interact with SlSBP13 at the protein level to interfere with *SlSBP13*-mediated *SFT* activation under LD conditions. To test this conjecture, we first conducted a transient transactivation assay in *N. benthamiana* leaves grown in LD using the *p35S::rSBP13* (a miR156-resistant *SlSBP13* version) and *p35S::SP5G* as effector constructs, along with the *pSFT::LUC*. As expected, luciferase activity significantly increased in the presence of the rSBP13, indicating activation of the *pSFT*. The presence of *SP5G* dampened *SlSBP13*-mediated activation, leading to reduced luciferase activity (Figure 5D). These findings suggested that SP5G antagonizes SlSBP13 transcriptional activity in the context of *SFT* regulation. We next asked whether SP5G and SlSBP13 interact at the protein level. Our data consistently supported the existence of an interaction between SP5G and SlSBP13, as confirmed by both Bimolecular fluorescence complementation (BiFC) and Fluorescence Resonance Energy Transfer (FRET) protein-protein interaction (PPI) assays (Figure 5E-5F; Figure S11A). Consistent with this interaction, Virus-Induced Gene Silencing (VIGS)-mediated silencing of *SlSBP13* enhanced the delayed flowering phenotype of *SP5G^pen^* plants grown under LD conditions (Figure S11B-S11C). Collectively, these findings suggest that high *SP5G* levels delay tomato flowering under LD conditions, at least in part, by altering the expression of specific miR156-targeted *SlSBPs* (Figure 2E; Table S1) and by repressing *SFT* via suppression of SlSBP13-mediated transcriptional activation.

### High gibberellin responses/levels enhance photoperiodic *SP5G*-mediated repression of tomato flowering

In *Arabidopsis*, DELLA proteins attenuate the transcriptional activity of CO, thereby reducing *FT* expression and delaying flowering (Xu et al., 2016; Wang et al., 2016). By contrast, the interaction between PRO/DELLA and SlCOL1 in tomato appears to be distinct, as *SP5G* transcript levels remained unchanged in *della*/*procera* (*pro*) mutant plants under LD conditions (Figure S9B), despite their reduced *SFT* expression and delayed flowering phenotype (Silva et al., 2019). Accordingly, the presence of a stable PROCERA version (PRO 17; Nir et al., 2017) did not significantly interfere with *COL1*-mediated activation of SP5G, as demonstrated by the luciferase transactivation assay (Figure S12). We then speculated that PRO/DELLA may modulate *SP5G* function at the protein level. Remarkably, BiFC assays revealed a physical interaction between PRO/DELLA and SP5G, but no significant interaction was detected by FRET (Figure S13). These observations indicated that the interaction between PRO/DELLA and SP5G is either very weak or absent. Thus, to better understand how the photoperiodic- and *PRO/DELLA*-dependent pathways interact at the genetic level, we introduced the *SP5G^pen^* allele into *pro* plants to generate *pro SP5G^pen^*. Regardless of photoperiod, *pro SP5G^pen^* plants exhibited a synergistic delay in flowering time compared to either parental line (Figure 6A-6C; Figure S14), despite showing no change in *SP5G* transcript levels (Figure S9B; Figure S15A). While time to anthesis in *pro* mutants was similar to MT, the presence of the *SP5G^pen^* allele in the *pro* background further delayed anthesis under both LD and SD conditions (Figure 6A–C; Figure S14). In addition, both *pro* and *pro SP5G^pen^* plants exhibited a stronger delay in flowering time under LD conditions (Figure S15B), suggesting a role for *PRO/DELLA* in photoperiodic responses. Nevertheless, *SP5G* and *PRO/DELLA* appear to act independently in the control of flowering time.

**Figure 6.**
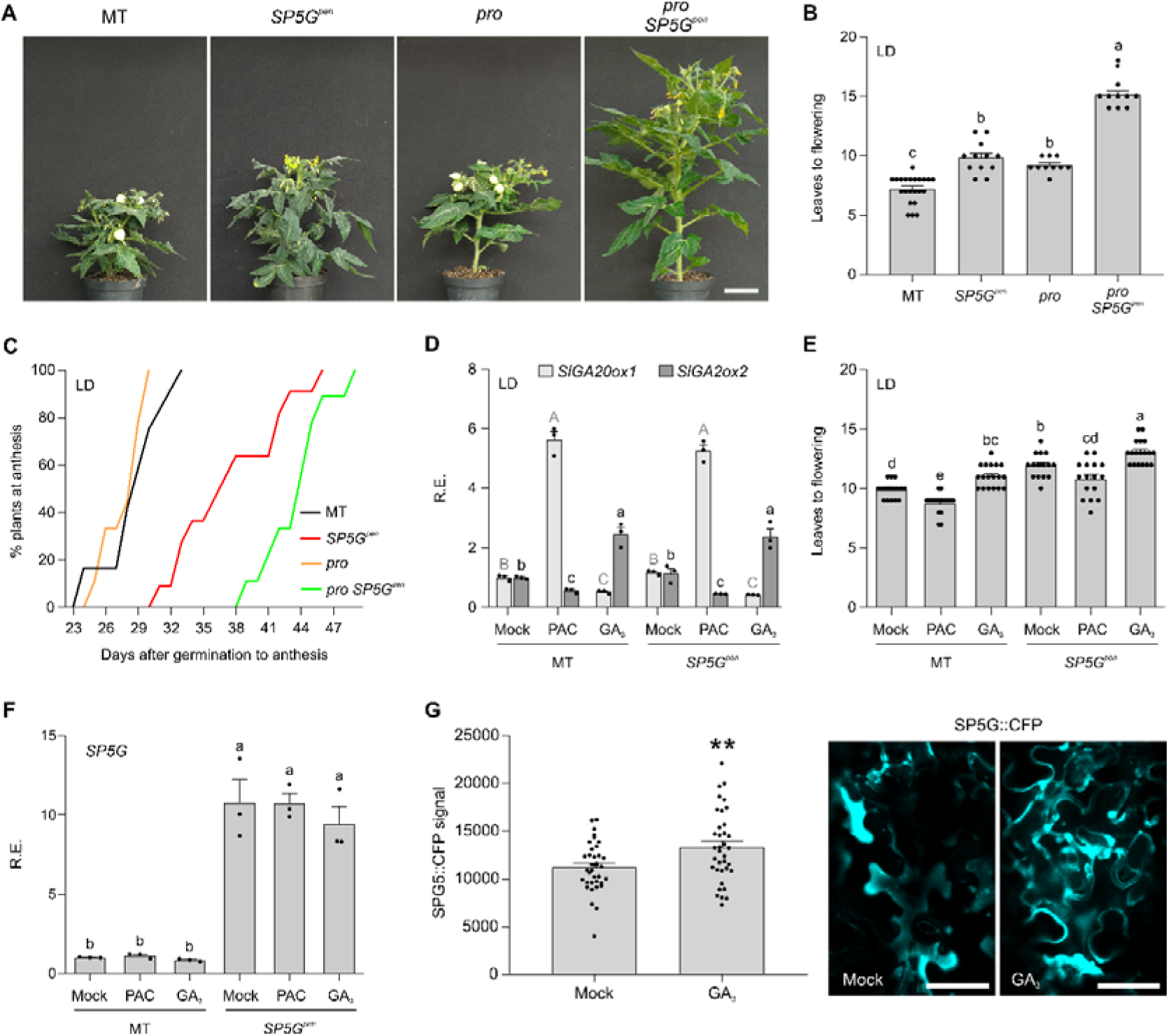
SP5G activity is enhanced by gibberellin (GA) in LD. **(A)** Representative images of 60-day-old MT, *SP5G^pen^*, *pro*, and *pro SP5G^pen^* plants cultivated in LD (14h/10h). Bar = 5cm. (**B and C)** Leaves to flowering (**B**) and percentage of plants at anthesis after germination in LD conditions (**C**) for MT, *SP5G^pen^*, *pro*, and *pro SP5G^pen^* plants. Data are expressed as mean ± SEM (Tukey’s test, *p<0.001*; n = 10-22). In (**C**), black, red, orange, and green lines represent MT, *SP5G^pen^*, *pro*, and *pro SP5G^pen^* genotypes, respectively. **(D)** Relative expression (R.E.) levels of *GA20ox1* and *GA2ox2* in 10-day-old MT and *SP5G^pen^* seedlings treated with mock, paclobutrazol (PAC), and gibberellin (GA_3_) in LD (ZT = 4; mean ± SEM; n = 3). Different uppercase and lowercase letters above the bars indicate significant differences in *SlGA20ox1* and *SlGA2ox2* expression among treatments, respectively (Tukey’s test, *p<0.05*). (**E)** Number of leaves to flowering of MT and *SP5G^pen^* plants treated with mock, PAC, and GA_3_ in LD (n = 15-18, mean ± SEM). Distinct letters indicate significant differences according to the Tukey’s test (*p<0.05*). **(F)** Relative expression (R.E.) levels of *SP5G* in 10-day-old MT and *SP5G^pen^* seedlings treated with mock, PAC, and GA_3_ in LD (ZT = 4; mean ± SEM; n = 3). Distinct letters indicate significant differences according to Tukey’s test (*p<0.05*). **(G)** Transient expression of the *p35S::SP5G::CFP* reporter in *N. benthamiana* leaves treated with mock and GA . Left: Quantification of CFP signal in leaf discs using a plate reader (Tecan Infinite 200 Pro, Switzerland). Values are mean ± SEM (Student’s t-test, *p < 0.01*, n = 36). Right: Representative fluorescence images of infiltrated *N. benthamiana* leaves after treatment. Scale bars = 50 µm. Each dot in all graphs represents individual data.

Interestingly, our RNA-seq analysis identified several DEGs related to GA biosynthesis and signalling, including some G*IBBERELLIN receptors*, G*IBBERELLIN-regulated proteins*, and *GIBBERELLIN 20-oxidase-1* and *-2* (*SlGA20ox1* and *SlGA20ox2*) genes (Figure 2E; Table S1–S4). These findings suggested altered GA levels or sensitivity in plants harbouring the *SP5G^pen^* allele. Thus, to investigate the direct effect of gibberellin on tomato photoperiodic flowering, we treated MT and *SP5G^pen^* seedlings after emergence (Fig. S20) with GA_3_ or the GA-biosynthesis inhibitor paclobutrazol (PAC) (Sun and Gubler, 2004; Silva et al., 2019). Consistent with the contrasting effects of GA_3_ and PAC on gibberellin homeostasis, *SlGA20ox1* and *SlGA2ox2* exhibited opposite expression patterns, reflecting feedback regulation of GA metabolism (Hedden and Thomas, 2012). In both genotypes, GA application reduced *SlGA20ox1* expression and promoted *SlGA2ox2*, whereas PAC treatment induced *SlGA20ox1* and reduced *SlGA2ox2* expression (Figure 6D). As expected, GA_3_-treated plants exhibited increased plant height, whereas PAC treatment resulted in plants with reduced height compared to mock treatment (Figure S15C, S20). Importantly, PAC-treated *SP5G^pen^* plants rescued MT flowering phenotype in LD, whereas GA_3_-treated MT displayed a flowering time similar to mock-treated *SP5G^pen^* plants (Figure 6E). In addition, GA_3_-treated *SP5G^pen^* plants mimicked the *pro SP5G^pen^* flowering phenotype in LD, displaying an increased number of leaves before flowering (Figure 6E).

Similar to the expression levels observed in the *pro* mutant background (Figure S9B; Figure S15A), *SP5G* transcripts remained unchanged in response to GA_3_ and PAC treatments compared to the mock control (Figure 6F), suggesting a potential post-transcriptional interaction between gibberellin and photoperiodic pathways. To investigate this possibility, we performed a transient assay in *N. benthamiana* leaves agroinfiltrated with the *p35S::SP5G::CFP* reporter construct and subsequently treated with GA_3_ or mock. Notably, GA_3_ treatment enhanced SP5G::CFP fluorescence (Figure 6G), indicating that gibberellin may indirectly modulate SP5G protein stability. Collectively, these findings support a model in which SP5G-mediated photoperiodic repression of flowering in tomato is at least partially regulated by gibberellin levels or signalling, particularly under LD conditions.

### Overlapping and unique contributions of *SP5G*, *PRO/DELLA*, and the miR156-*SlSBP* module to the regulation of *SFT-*dependent and *-*independent flowering pathways

*SFT* expression is modulated by components of the photoperiodic, age-dependent, and gibberellin pathways (Figure S16; Soyk et al., 2017; Silva et al., 2019; Cui et al., 2020). To further explore the interactions between these pathways and *SFT* at the genetic level, we crossed *SP5G^pen^*, *miR156OE*, and *pro* each with the *sft* mutant in the MT background (Vicente et al., 2015), generating the *sft SP5G^pen^*, *sft miR156OE*, and *sft pro* genotypes. Plants harbouring either the *miRNA156OE* construct or *SP5G^pen^*allele in the *sft* background exhibited a synergistic effect on delaying flowering time, independent of the photoperiodic condition. In LD, *sft SP5G^pen^* plants exhibited the most pronounced flowering delay, both in terms of leaves to flowering and days to anthesis (Figure 7A-7C), indicating that SP5G does not act exclusively via *SFT*. In SD, *sft SP5G^pen^* plants also showed a longer time to anthesis, but the highest number of leaves to first inflorescence was observed in *sft miR156OE* plants (Figure S17A-S17C). Strikingly, we observed no difference in the number of leaves to flowering between *sft* and *sft pro* plants (Figure 7A-7C; Figure S17A-S17C). Nevertheless, *sft* plants harbouring either the loss-of-function *pro/della* allele or the *miR156OE* construct exhibited earlier anthesis (Figure 7A-7C; Figure S17A-S17C). Collectively, our data support the notion that PROCERA/DELLA primarily regulates tomato floral transition via *SFT*, while the miR156–*SlSBP* module and *SP5G* may also modulate *SFT-* independent flowering pathways.

**Figure 7.**
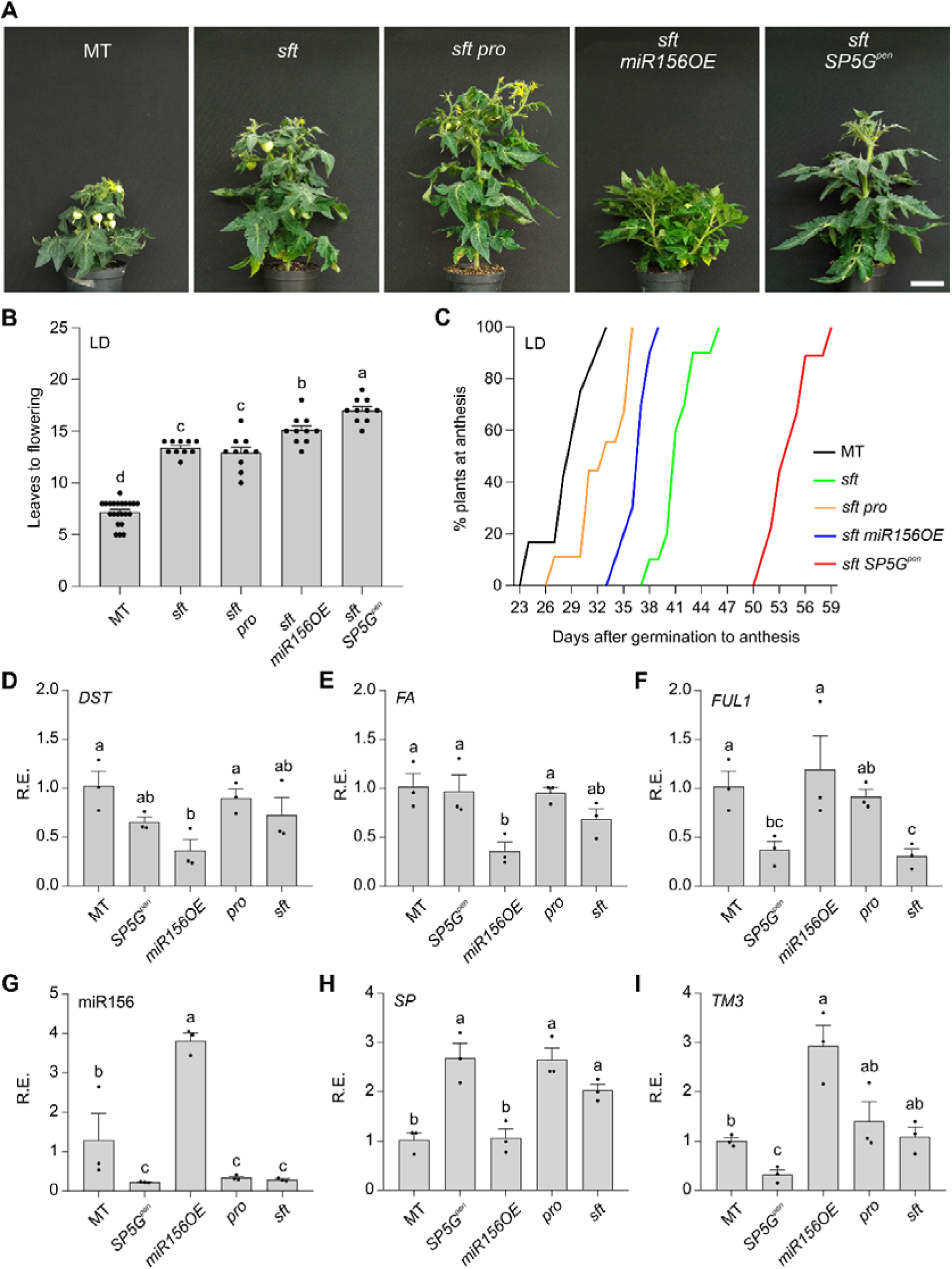
Photoperiod, age, and gibberellin regulate flowering in an *SFT*-dependent and independent manner. **(A)** Representative images of 60-day-old MT, *sft*, *sft pro*, *sft miR156OE*, and *sft SP5G^pen^* plants cultivated in LD (14h/10h). Bar = 5cm. (**B, C)** Leaves to flowering (**B**) and percentage of plants at anthesis after germination in LD conditions (**C**) for MT, *sft*, *sft pro*, *sft miR156OE*, and *sft SP5G^pen^* plants. Data are expressed as mean ± SEM (n = 10-22). (**D-I)** Relative expression (R.E.) of *DST* (**D**), *FA* (**E**), *FUL1* (**F**), miR156 (**G**), *SP* (**H**), and *TM3* (**I**) in MT, *SP5G^pen^*, *miR156OE*, *pro*, and *sft* vegetative apices in LD (ZT = 4; mean ± SEM; n = 3). Distinct letters indicate significant differences according to Tukey’s test (*p<0.05*). Each dot represents individual data.

It was recently reported that *DELAYED SYMPODIAL TERMINATION* (*DST*) and *SFT* promote tomato flowering by regulating shared and specific developmental marker genes for shoot apical meristem (SAM) maturation (Meir et al., 2021). We then asked how *SP5G*, *PRO/DELLA*, and the miR156-targeted *SlSBPs* contribute to the regulation of these developmental markers at the SAM. To that end, we dissected shoot apices at the vegetative stage (including the SAM and two leaf primordia; Figure S18) from MT, *SP5G^pen^*, miR156OE, *pro*, and *sft* seedlings grown under LD conditions, and quantified the transcript levels of flowering-related genes by qRT-PCR. These genes were selected based on our RNA-seq analysis (Table S1-S4) and from RNA-seq data mined from Meir et al. (2021). Our results showed that *DST* and *DST*-dependent *FALSIFLORA (FA,* a *LEAFY* homologue, Molinero-Rosales et al., 1999) transcript levels were reduced only in shoot apices of *miR156OE*, whereas *FRUTFULL1* (*FUL1*), an *SFT*-dependent meristem marker (Meir et al., 2021), was downregulated in *sft* and *SP5G^pen^* apices (Figure 7D-7F). In contrast, *FUL2* expression did not change across the genotypes (Figure S17D). miR156-targeted *SlSBP3a*, *SlSBP4b* and *SlSBP15* were downregulated in the shoot apices of *miR156OE* seedlings, whereas *SlSBP3a* and *SlSBP4b* were upregulated in *SP5G^pen^*, *pro*, and *sft* genotypes (Figure S17E-S17G), consistent with the lower levels of mature miR156 observed in these lines (Figure 7G). *SlSBP15* and *SlSBP3a* transcripts have been reported to accumulate at higher levels in the apices of both *sft* and *dst* mutants, likely due to a compensatory mechanism activated in response to impaired floral transition (Meir et al., 2021). Interestingly, shoot apices of *SP5G^pen^*, *pro*, and *sft* mutants also displayed elevated transcript levels of *SELF-PRUNING* (*SP*) (Figure 7H), a well-characterized antiflorigen that plays a central role in maintaining the sympodial growth habit of tomato (Lifschitz et al., 2014).

Our data suggested that *DST* and *FA* (Meir et al., 2021) act in the shoot apex primarily downstream of the miR156–*SlSBP* module, whereas *FUL1* may function downstream of the *SP5G*-, *PRO/DELLA*-, and *SFT*-dependent pathways. We then quantified the transcript levels of *DST*, *FA*, and *FUL1* (Meir et al., 2021) in the shoot apices of *sft SP5G^pen^*, *sft miR156OE*, and *sft pro* seedlings. The expression of *DST* and *FA* was reduced only in the miR156-overexpressing *sft* shoot apices. *FUL1* was downregulated in the *sft* apices harbouring the *pro* and *SP5G^pen^* alleles, but upregulated in *sft miR156OE* (Figure S17H-S17J). Thus, our data reinforced the notion that the age-dependent pathway modulates both *SFT*- and *DST*-dependent floral transitions, while *PRO/DELLA* and *SP5G* act primarily through the *SFT*-dependent pathway in the shoot apices. However, the *sft SP5G^pen^* double mutant exhibited a more pronounced delay in flowering compared with *sft* (Figure 7A–C; Figure S17A–S17C), suggesting the involvement of an additional flowering pathway in this genotype. In this context, *TOMATO MADS 3 (TM3)* emerged as a strong candidate, since both *SP5G^pen^* and *sft SP5G^pen^* displayed reduced *TM3* expression at the SAM (Figure 7I; Figure S17K). In addition, no significant changes in *SFT* or *FA* expression were detected in the vegetative meristems of the late-flowering *tm3 stm3* (*STM3*, *SISTER OF TM3*) mutant (Zahn et al., 2023). Taken together, these observations suggested that *SP5G^pen^* may regulate the floral transition at the SAM independently of *SFT* by modulating *TM3* expression.

## DISCUSSION

The transition from short-day to day-neutral flowering during tomato domestication was largely driven by mutations in the *SP5G* locus, which substantially reduced photoperiodic sensitivity (Soyk et al., 2017; Zhang et al., 2018). However, the mechanisms regulating *SP5G* activity under long-day conditions and its integration with other flowering pathways remained unclear. Here, we uncovered an additional regulatory layer of photoperiodic flowering in tomato and wild relatives by analysing CRISPR mutants and a photoperiod-sensitive tomato line carrying a wild *SP5G* allele. Our data show that tomato cv. MT harbouring the *SP5G* allele from *S. pennellii* exhibited late flowering in LD (Figure S1). Importantly, we revealed that *COL1* modulates flowering time in tomato and *S. pimpinellifolium*, and that COL1 directly activates *SP5G* in LD and enhances its repressive role on flowering (Figure S2; Figure 3). The SlCOL1-based *SP5G* activation indicates a conserved role of CO in controlling the photoperiod flowering program. However, while CO promotes flowering in *Arabidopsis* by directly activating *FT* under LD conditions (Robson et al., 2001; An et al., 2004; Tiwari et al., 2010), SlCOL1 delays flowering in tomato by activating the floral repressor *SP5G* (Figure 2 and 3), and repressing *SFT* (Cui et al., 2022). Our data suggested that additional cofactors or regulatory elements may be necessary to modulate *SP5G* expression (Figure 3; Zhang et al., 2018). In *Arabidopsis*, CO alone exhibits low binding affinity to the *FT* CORE element and requires the NF-YB2/NF-YC3 complex for effective binding (Gnesutta et al., 2017). Further studies will determine whether this is also the case for SlCOL1-dependent *SP5G* direct activation.

CO protein stability is modulated by phytochromes, which impacts the activity of its downstream targets (Valverde, 2004; Shim et al., 2017; Luo et al., 2022; Romero et al., 2024). In *Arabidopsis*, whilst PHYA stabilizes CO in the evening to promote *FT* transcription, PHYB induces CO degradation in the morning (Imaizumi and Kay, 2006; Valverde, 2011; Shim et al., 2017; Luo et al., 2022). Conversely, in potato, StPHYB stabilizes StCOL1 in the morning, leading to *StSP5G* activation (Abelenda et al., 2016). Our genetic and molecular analyses strongly indicate that SlCOL1 protein activity also depends on PHYB1, as the *phyB1* mutation suppresses the effects of *SlCOL1* overexpression. Thus, we speculate that PHYB1 may also modulate SlCOL1 activity post-transcriptionally, possibly by affecting its protein level.

Although *SP5G* transcription is drastically reduced in the absence of functional *PHYB1* (Figure 4B), *PHYA* and *PHYB2* appear to have no significant impact on tomato photoperiodic flowering (Cao et al., 2016; 2018). Here, we showed that the *phyB1* mutation is sufficient to rescue vegetative and late flowering phenotypes of plants carrying either the *SlCOL1*-overexpressing construct or the *SP5G^pen^* allele (Figure 4; Figure S7). These findings support the notion that *PHYB1* is the key photoreceptor integrating environmental signals to regulate flowering time and vegetative architecture. Our results are in line with findings in potato, where *StPHYB*, *StCOL1*, and *StSP5G* modulate tuber induction through similar regulatory mechanisms (Abelenda et al. 2016). This conserved molecular circuit supports the existence of an evolutionary link between flowering and tuberization within the Solanaceae family (Jing et al., 2023). Notably, it was recently reported that tuberization in cultivated potato arose from an ancient hybridization between tomato and Etuberosum ancestors (Zhang et al., 2025).

We previously reported that gibberellin and the miR156-*SlSBP* module are interconnected in tomato through the regulation of *SFT, SlSBPs,* and flower identity genes (Silva et al., 2019). Similar to what was reported in *Arabidopsis* (Jung et al., 2012; 2016), we show here that the miR156–*SlSBP* module acts primarily downstream of the photoperiodic pathway, as high levels of *SlCOL1* and *SP5G* repress flowering at least in part by modifying *SlSBP* transcript levels in seedlings (Figure 2; Table S1-S4). However, in tomato, the photoperiod and age-dependent pathway also converge to regulate flowering under LD conditions at the protein level through the repressive role of SP5G on SlSBP13-dependent *SFT* activation. We propose that SP5G may act as a transcriptional cofactor, similar to its *Arabidopsis* homologs FT and TERMINAL FLOWER1 or TFL1 (Ho and Weigel, 2020). This likely explains the more pronounced delay in flowering time observed in the *miR156OE SP5G^pen^* plants in LD when compared with *miR156OE* or *SP5G^pen^* plants growing under similar conditions (Figure 5).

In *Arabidopsis*, CO transcriptional activity is attenuated by DELLA proteins to prevent *FT* transcription (Xu et al., 2016; Wang et al., 2016). In tomato, elevated gibberellin levels/responses delay flowering by reducing *SFT* expression, an effect that is partially counteracted by strigolactone (SL) treatment under LD conditions (Silva et al, 2019; Visentin et al., 2024). Here, we showed that tomato plants exhibited increased sensitivity to photoperiod under enhanced GA responses, and reduced GA levels were sufficient to rescue the late flowering phenotype of *SP5G^pen^* plants (Figure 6E). These results are consistent with previous findings in rice, an SD plant, in which cultivars with reduced GA sensitivity also show impaired photoperiod responsiveness (Evans, 1993). Importantly, our transcriptomic data suggested that the *SlCOL1*–*SP5G*-dependent photoperiodic circuit modulates GA responses potentially through GA signalling and homeostasis (Figure 2E; Table S1-S4). Although our genetic and molecular data indicated that *PRO/DELLA* functions largely independently of the *SlCOL1*–*SP5G*-regulated photoperiodic pathway (Figure 6; Figure S9; Figure S12-S15), additional photoperiod- and light-dependent flowering activators, such as SlPIF4, physically interact with PRO/DELLA to activate *SFT* and promote tomato flowering time and inflorescence development (Moreira et al., 2025). Notably, exogenous GA application did not alter *SP5G* transcript levels but instead affected SP5G protein activity, suggesting a post-transcriptional interaction between gibberellin and photoperiodic pathways (Figures 6F and 6G). Although the underlying mechanisms remain unclear, this interaction may involve post-translational modifications, such as phosphorylation or ubiquitination, or chaperone-mediated regulation of protein turnover.

Tomato floral transition at the SAM is primarily orchestrated by two main pathways: one systemic and another local (autonomous), regulated by the *SFT* and *DST* genes, respectively (Meir et al., 2021). Our results corroborated previous findings that *PRO/DELLA* does not regulate *FA* activity in the shoot apices (Figure 7E; Silva et al., 2019), which indicated that *PRO/DELLA* functions primarily through the *SFT*-dependent systemic pathway (Meir et al., 2021), as further supported by the flowering time phenotype of the double *sft pro* mutant (Figure 7A-7C; Figure S17A-S17C). Our genetic and molecular analyses indicated that the miR156-targeted *SlSBPs* act through both systemic and autonomous pathways, regulating *SFT* transcription in leaves and *FUL1*, *DST*, and *FA* at the SAM (Figure 7D-7F; Figure S16; Silva et. al., 2019). Similarly, *SP5G* seems to regulate flowering via both *SFT*-dependent and independent pathways, as evidenced by the phenotype of the double *sft SP5G^pen^* mutant and *TM3* regulation at the shoot apex (Figure 7A-7C, 7I; Figure S17k). *PRO/DELLA* plays a repressive role in inflorescence development, as *sft pro* plants presented a similar number of leaves to *sft*, but their anthesis occurred earlier. Consistent with this, the increased GA sensitivity observed in plants overexpressing miR156 (Miao et al., 2019; Ferigolo et al., 2023) may account for the earlier anthesis in *sft miR156OE*, despite their production of more leaves to flowering compared to *sft* (Figure 7B, 7C). These results support a dichotomous role of the *PRO/DELLA*-regulated flowering pathway in tomato, acting as a promoter during the floral transition but as a repressor of inflorescence development. In contrast, *SP5G* functions as a negative regulator of both processes.

In summary, our findings expand the current understanding of tomato flowering by identifying *SlCOL1* as a key regulator of the photoperiodic response, acting through the transcriptional activation of *SP5G*. *SP5G*, in turn, interplays synergistically with the age-dependent and gibberellin pathways at multiple regulatory levels (Figure 8). Given the established role of SBP/SPL transcription factors in promoting flowering (Silva et al., 2019; Cui et al., 2020), the SP5G interaction with SlSBP13 suggests a mechanism whereby SP5G sequesters SlSBP13, impairing its activation of SFT and delaying the floral transition. These integrated genetic insights provide a framework for fine-tuning the balance between vegetative and reproductive growth in tomato (Vicente et al., 2015), facilitating the development of higher-yielding cultivars and more sustainable agricultural practices in the coming decades.

**Figure 8.**
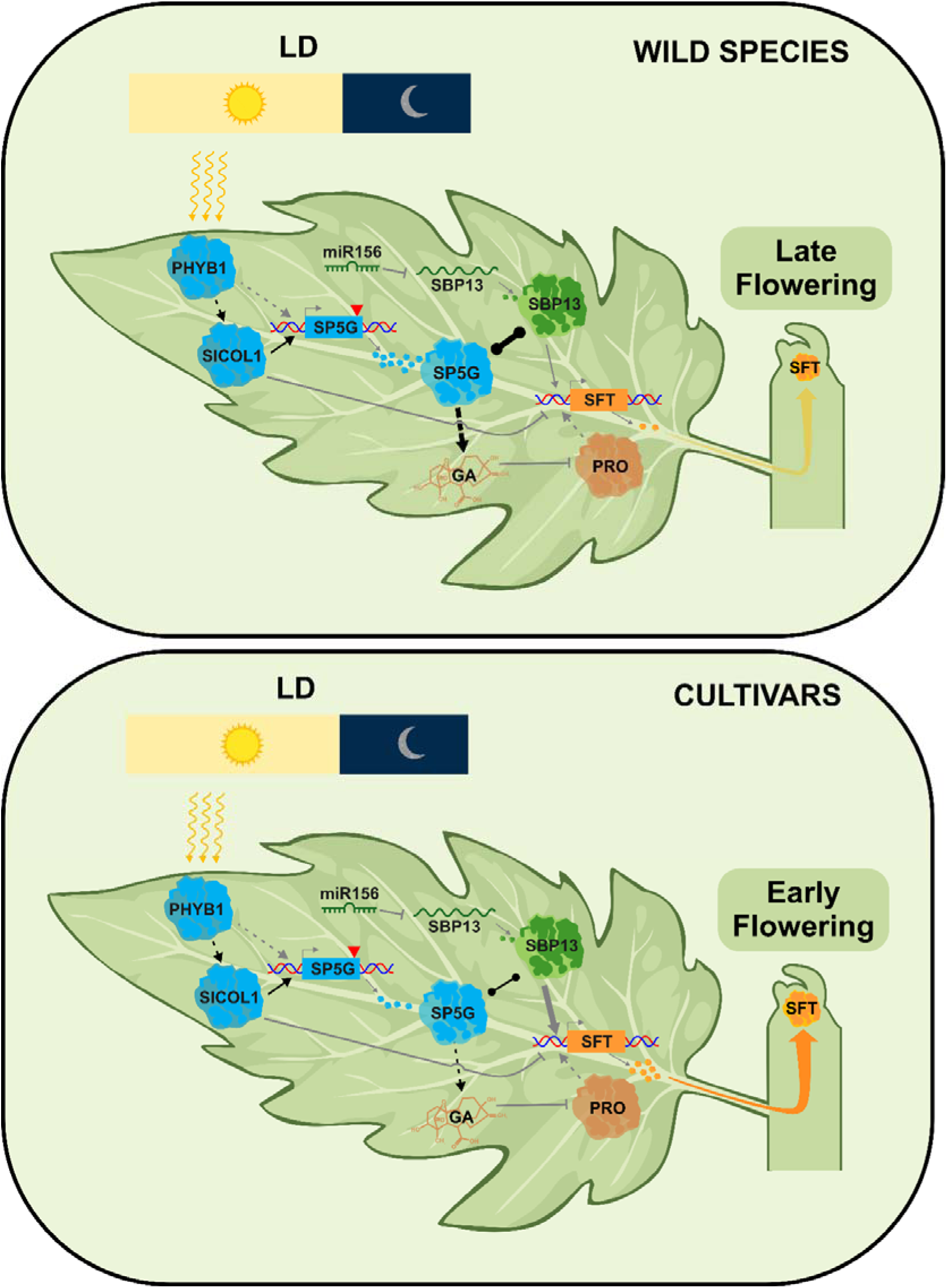
Working model for flowering control under long-day (LD) conditions in wild and cultivated tomatoes. Under LD, the photoreceptor PHYB1 modulates SlCOL1 activity during the early hours of the day, promoting *SP5G* transcription. SP5G, in turn, represses *SFT* expression in mature leaves by interfering with the miR156–*SlSBP13* activation module. The gibberellin (GA) repressor PROCERA/DELLA promotes *SFT* expression, while GA levels and sensitivity are photoperiod-dependent. Once produced in the leaves, SFT moves to the shoot apex to trigger flowering. In wild species, stronger SP5G activity enhances GA sensitivity and restricts SlSBP13-mediated *SFT* activation, delaying flowering. By contrast, reduced SP5G activity in cultivated tomato allows SlSBP13 and PRO/DELLA to promote *SFT* expression, resulting in earlier flowering. Arrows represent transcriptional activation; T-shaped lines indicate repression; lines with circles denote protein–protein interactions. Solid lines indicate direct regulation, and dashed lines indicate indirect or unconfirmed regulation. Black lines highlight regulatory relationships identified in this study.

## MATERIALS AND METHODS

### Plant material and growth conditions

Plants used in this work were in the tomato cv. Micro-Tom (MT, LA3911) background. *pro*, *miR156OE*, and *sft* were described elsewhere (Vicente et al., 2015; Silva et al., 2019). The *phyB1* and *SP5G^pen^* plants were generated through introgression from the *temporarily red light-insensitive* (*tri*; LA3808) mutant and the IL 5-4, respectively, into the MT background as previously described (Carvalho et al., 2011). Seeds of *S. pimpinellifolium* (LA1584) were kindly donated by the Tomato Genetics Resource Center (TGRC, Davis, University of California). The accession LA1584 was selected due to its stronger photoperiodic flowering responses (Soyk et al., 2017). Plants were grown as described in Silva et al. (2019). Genotypes carrying double mutations and/or transgenes were generated through crosses between individual genotypes, and presence of mutations or transgenes was confirmed by PCR analysis and sequencing.

Photoperiod-related experiments involving tomato, *S. pimpinellifolium*, and *N. benthamiana* were conducted in controlled-environment growth chambers. Conditions were set at 27 °C, with PAR irradiance of 200 µmol m□□² s□□¹, and relative humidity maintained between 40–60%. Plants were grown under either long-day (14 h light/10 h dark) or short-day (10 h light/14 h dark) photoperiods. The genotypes used in this study are listed in Table S5.

### Genotyping and Phenotyping

Genomic DNA was extracted from young leaves as described by Fulton et al. (1995). PCR amplification was carried out using GoTaq® Green Master Mix (Promega). When necessary, restriction enzyme digestion was performed to distinguish mutant alleles, according to the manufacturer’s instructions (New England Biolabs, Bethesda, MD, USA). PCR products were visualized on 1.5% (w/v) agarose gels. The primers and restriction enzymes used for genotyping are listed in Table S6.

Flowering time was characterized by measuring the number of leaves produced before the first inflorescence, and the number of days to anthesis of the first flower, which integrates the timing of shoot apex transition with endogenous and environmental factors that affect growth rate and floral bud development (Soyk et al., 2017). At least eight plants/genotype were used for each analysis.

### RNA extraction, cDNA synthesis, and qRT–PCR analysis

Total RNA was extracted from vegetative apices, mature leaves, or seedlings using TRIZOL® reagent (Invitrogen), following the manufacturer’s instructions. After extraction, samples were treated with DNase I (Thermo Fisher Scientific) to remove genomic DNA. cDNA synthesis was performed using the pulsed stem-loop method, which enables the detection of small mature RNAs such as miR156 (Varkonyi-Gasic et al., 2007), from 1 µg of RNA total. Quantitative reverse transcription–PCRs (qRT– PCRs) were performed using Platinum SYBR Green qPCR SuperMix UDG (Invitrogen) and analysed on a Step-OnePlus real-time PCR system (Applied Biosystems). *ACTIN4* was selected as an internal control after initially analysing three housekeeping genes: *ACTIN4*, *BETA-TUBULIN 1*, and *ALPHA-TUBULIN 3*. For each genotype, at least three biological and two technical replicates were used in the qRT– PCR analyses. Relative gene expression (RE)s was calculated using the 2^−ΔΔCT^ method (Livak and Schmittgen, 2001). The primers used are listed in Table S6.

### Vector constructs and plant transformation

To generate the *p35S::SlCOL1* construct used for plant transformation, the full-length coding sequences (CDS) of *SlCOL1* were cloned into the *pENTR D-TOPO* entry vector (Invitrogen) and subsequently recombined into the destination vector *pK7WG2.0* using LR Clonase (Thermo Fisher Scientific), through the Gateway cloning system (Karimi et al., 2002). Plant transformation was performed by Pangeia Biotech (Unicamp, Campinas, SP, Brazil) using *Agrobacterium tumefaciens* strain LBA4404. At least six independent *p35S::SlCOL1* transgenic events were obtained, of which three were selected for further analysis.

Constructs for transactivation assays were generated as follows: a miR156-resistant version of *SlSBP13*, referred here as *rSBP13*, was produced by introducing PCR-mediated synonymous mutations into the miR156-binding site. Full-length CDSs of *rSBP13* and *SP5G* were cloned into the universal GoldenBraid acceptor vector pUPD2 (Vazquez-Vilar et al., 2017), and entry vectors containing both CDSs flanked by attL1 and attL2 sites were subsequently generated using the BIG cloning system (Ferigolo et al., 2022). The *SP5G* CDS was cloned both with and without a stop codon. In addition, the universal GoldenBraid system was used to generate *p35S*::*rSBP13* and *p35S*::*SP5G* constructs in the Golden Gate acceptor vector pICSL86900OD (Vazquez-Vilar et al., 2017). The PRO CDS was previously cloned into *pENTR D-TOPO* by Silva et al. (2019).

The *pSP5G^lyc^*::*LUC*::*3’UTR^lyc^* and *pSP5G^lyc^*::*LUC*::*3’UTR^pimp^* constructs were produced by modifying the pGreen_dualluc_*3*′*UTR*_sensor vector, which contains the Cauliflower Mosaic Virus (CaMV) 35S promoter driving the expression of firefly luciferase (*p35S::LUC*). Briefly, the *p35S::LUC* sequence was replaced with *pSP5G^lyc^::LUC* using PCR complementation, restriction enzymes, and T4 DNA ligase (Promega). The *pSP5G^lyc^* and *LUC* fragments were cloned individually and fused using PCR complementation. After generating a pGreen_dualluc_*3*′*UTR*_sensor vector harbouring the *pSP5G::LUC* sequence, the 3′UTRs from *S. lycopersicum* (MT, *3*′*UTR^lyc^*) and *S. pimpinellifolium* (LA1589, *3*′*UTR^pimp^*) were cloned to produce the constructs *pSP5G::LUC::3*′*UTR^lyc^* and *pSP5G::LUC-3*′*UTR^pimp^*. The *SFT* promoter was cloned into the pGreenII 0800-LUC vector using restriction enzymes and T4 DNA ligase. Promoter fragments of 1994 bp (*SP5G*) and 1006 bp (*SFT*), upstream of their respective start codons, were amplified using genomic DNA as templates. The 3′UTRs were approximately 450 bp in length for both constructs. Primers used for all constructs are listed in Table S6.

### Protein isolation and western blotting

Proteins were isolated from seedlings at 10 post-germination days (DPG) using the TRIZOL® reagent (Invitrogen), following the manufacturer’s instructions. Protein concentrations were determined by Bradford assays (Bio-Rad), using albumin as the standard and following the manufacturer’s protocol. Proteins were then separated by SDS-PAGE using standard procedures, transferred to PVDF membranes (Bio-Rad), and probed with AtCO antibody (αAtCO; Valverde et al., 2004). Membranes were subsequently incubated with a goat anti-rabbit IgG secondary antibody (Thermo Fisher Scientific). Blots were developed using the Clarity™ Western ECL Substrate (Bio-Rad), according to the manufacturer’s instructions, and analysed using an Amersham Imager 680 (GE Healthcare). Membranes were stained with Ponceau Red to verify equal protein loading.

### Clustered regularly interspaced palindromic repeats (CRISPR)/CRISPR-associated protein 9 (Cas9) gene editing

*SlCOL1* was targeted with two single-guide RNAs (sgRNAs) for CRISPR/Cas9-induced gene editing. Protospacer sequences were selected using CRISPR/Direct and CCTop— CRISPR/Cas9 target online predictor tools (Naito et al., 2015; Stemmer et al., 2015).

The *pDIRECT_22C* vector (Čermák et al., 2017, Addgene plasmid # 91135) was used as a template to assemble spacers, scaffolds, and the P2A/Csy4 splicing systems under the control of the *CmYLCV* promoter. PCR products and vector assembly were generated as previously described for the *pDIRECT_22C* vector (Čermák et al., 2017).

Due to the high sequence similarity between *SlCOL1* and its ortholog in *S. pimpinellifolium*, the *SlCOL1* sgRNAs also targeted *SpCOL1*. *A. tumefaciens* strain LBA4404 harbouring the *pDIRECT_22C* vector with *SlCOL1* sgRNAs was used to transform tomato (cv. MT) and *S. pimpinellifolium* (LA1584), following the protocol described by Pino et al. (2010). T0 plants were first genotyped by PCR to confirm the presence of *Cas9*, followed by PCR amplification using primers flanking the sgRNA target sites to detect deletions in the *SlCOL1/SpCOL1* locus. Mutations were confirmed by Sanger sequencing. The primers used are listed in Table S6.

### RNA-seq

Total RNA was extracted from two-week-old seedlings grown under LD conditions using the E.Z.N.A. Plant RNA Kit (Omega), following the manufacturer’s instructions. Two independent biological replicates from each genotype were used for RNA-seq library construction and high-throughput sequencing (Illumina NovaSeq platform) at CABIMER (Andalusian Molecular Biology and Regenerative Medicine Centre, Spain). Samples generated approximately 30 million single-end reads each. Adapter sequences and low-quality reads were removed from the raw data. The resulting high-quality reads were mapped to the tomato reference genome ITAG4.0 (Tomato Genome Consortium, 2012) using HISAT2 (Kim et al., 2019). StringTie (Kovaka et al., 2019) and the Bioconductor package ballgown (Frazee et al., 2015) were employed to assemble and quantify transcripts with default parameters, respectively. The reproducibility between replicates was evaluated by Principal Component Analysis (Figure S19).

Differential gene expression (DEG) analysis was performed using the Bioconductor R package limma v.3.444 (Ritchie et al., 2015). DEGs from *SP5G^pen^*, *p35S::SlCOL1*, and *p35S::SlCOL1 SP5G^pen^* seedlings compared with MT, with a *p-value* ≤0.05 and fold-change ≤0.7 or ≥ 1.5, were annotated based on the SOL Genomics Network (version SL4.0, annotation ITAG4.0). Venn diagrams were generated online at https://www.interactivenn.net/. Shared DEGs among these genotypes related to photoperiodic, age, and GA flowering pathways, as well as flowering markers described by Meir et al. (2021), were selected, and a heatmap was generated using the Heatmapper online tool (Babicki et al., 2016).

### Luciferase transactivation assays

For transactivation assays, reporter and effector constructs were agroinfiltrated into leaves of 5-week-old *N. benthamiana* plants. After three days, leaves were sprayed with D-luciferin (25 mg/L) (Promega), and LUC activity was evaluated using the NEWTON 7.0 CCD imaging system (Vilber). Relative LUC activity was quantified using the NEWTON 7.0 software (https://www.vilber.com/newton-7-0/) with default settings. For *SP5G* promoter activity analysis, *pSP5G::LUC::3*′*UTR^lyc^* and *pSP5G::LUC-3*′*UTR^pimp^*reporter constructs were co-infiltrated with either *p35S::SlCOL1* or *p35S::NLS-GFP* effector constructs. The *pSFT::LUC* reporter was co-infiltrated with *p35S::rSBP13* plus *p35S::NLS-GFP*, *p35S::SP5G* plus *p35S::NLS-GFP*, or *p35S::NLS-GFP* alone. To ensure comparability, the final bacterial concentration was standardized across all infiltration mixtures (OD = 0.6). *p35S::NLS-GFP* construct was used as a negative control (Barrera-Rojas et al., 2023).

### Electrophoretic mobility shift assays (EMSAs)

EMSAs were performed with minor modifications to the protocol described by Smaczniak et al. (2012). *SlCOL1* was PCR-amplified from the cDNA of *S. lycopersicum* cv. Moneyberg and cloned into the *pSPUTK* vector (Stratagene). In vitro protein synthesis was carried out using the TnT SP6 High-Yield Wheat Germ Protein Expression System (Promega), following the manufacturer’s instructions. DNA probes were amplified from genomic DNA and cloned into the *pJET* vector (Thermo Scientific). Fluorescent labelling was performed via PCR using DY-682 labelled primers specific to the *pJET* backbone. Labelled probes were purified with the NucleoSpin Gel and PCR Clean-up Kit (Macherey-Nagel). Binding reactions were assembled by incubating the in vitro–synthesized proteins with labelled probes, and protein–DNA complexes were resolved on native polyacrylamide gels. Gels were scanned and imaged using a Li-Cor Odyssey system. The primers used are listed in Table S6.

### Chromatin immunoprecipitation (ChIP)-qPCR

ChIP assay was performed on 10-DAG *p35S::SlCOL1* seedlings growing under LD conditions using the EpiQuik Plant ChIP kit (Epigentek Group, USA), following the manufacturer’s instructions. Briefly, 10 g of tissues were cross-linked with 1.0% (v/v) formaldehyde solution through vacuum infiltration for 15 minutes. Nuclei were then isolated, and chromatin was fragmented by sonication to obtain DNA fragments ranging between 0.3 – 1 kb. Chromatin fragments were immunoprecipitated using either the anti-AtCO antibody (Valverde et al., 2004) or a non-immune IgG antibody (αIgG) provided by the EpiQuik Plant ChIP kit (Epigentek Group, USA). Enrichment of SlCOL1-bound DNA regions was evaluated by qPCR using the primers listed in Table S6. Fold enrichment was calculated by normalizing αAtCO values to those obtained with a non-immune antibody control for each sample. Three biological and three technical replicates were analysed for the enrichment of each tested region.

### BiFC and FRET assays

BiFC experiments were performed as described by Ferigolo et al. (2022). In brief, the CDSs of rSBP13 and PRO were fused to the N-terminal of Yellow Fluorescent Protein (YFP) in the pSITE-BiFC-nEYFP-C1 vector, while SP5G CDS was fused to the C-terminal region of YFP in the pSITE-BiFC-cEYFP-C1 vector. *A. tumefaciens* strain GV3101 carrying the respective constructs was co-infiltrated into *N. benthamiana* leaves. After 48h, leaf discs were analysed using a confocal microscope (3500 Genetic Analyzer, Applied Biosystems). Empty vector combinations were used as negative controls. *AtWWP1::mCherry* was co-expressed as a nuclear marker (Silva et al., 2015). Agrobacteria harbouring a binary plasmid encoding p19 protein from Tomato Bushy Stunt Virus (TBSV), a suppressor of post-transcriptional gene silencing (Qiu et al., 2002), were also co-infiltrated.

For FRET experiments, the *SP5G* CDS lacking the stop codon was cloned into pGWB644, while the CDSs of *rSBP13* and *PRO*Δ*17* were cloned into pGWB642 using LR Clonase (Thermo Fisher Scientific). The resulting SP5G::CFP construct was then co-infiltrated with either YFP::rSBP13 or YFP::PRO into *N. benthamiana* leaves via agroinfiltration. As a negative control, the empty pGWB642 vector was co-infiltrated with *SP5G::CFP*. Three days after transfection, epidermal cells were visualized using a confocal laser scanning microscope (Olympus FV3000). CFP was excited with a 455-nm laser, and YFP was excited with a 514-nm laser. FRET was measured by the acceptor photobleaching method (Ishikawa-Ankerhold et al., 2012). Briefly, a three-second pulse of high-intensity laser at 514 nm was applied to bleach the YFP signal acceptor in a whole cell. Pre-bleaching and post-bleaching CFP (donor) fluorescence intensities were recorded. FRET efficiency (E_FRET_) was calculated using the equation: E_FRET_ = 1-FDA/FD, where FDA and FD indicate CFP signal before and after photobleaching, respectively.

### VIGS assay

The VIGS vectors TRV1 and TRV2-LIC (pYL170) were described previously (Dong et al., 2007). *TRV2-SlSBP13* and *pTRV2-SlPDS* were generated following the procedure described by Dong et al. (2007). Briefly, target fragments were PCR-amplified from MT leaf cDNA using primers listed in Table S6. Purified PCR products (100 ng) were treated with T4 DNA polymerase (New England Biolabs) in the presence of 5 mM dATP at 22 °C for 30 min, followed by enzyme inactivation at 70 °C for 20 min. The TRV2-LIC vector was digested with PstI restriction enzyme and treated with T4 DNA polymerase similarly, except that dTTP replaced dATP. Equal amounts (50 ng) of the treated insert and vector were mixed, incubated at 65 °C for 2 min and 22 °C for 10 min, and 10 µL of the reaction was transformed into *E. coli* TOP10. TRV1, TRV2-*SlSBP13*, and TRV2-*SlPDS* were then introduced into *A. tumefaciens* GV3101.

For VIGS, five-day-old *SP5G^pen^* seedlings were vacuum-infiltrated with a mixture of TRV1 and either TRV2-*SlSBP13* or TRV2-*SlPDS.* Infiltration was performed at 760 mmHg for 5 min, and pressure was slowly released thereafter. Seedlings were gently washed three times with sterile distilled water and transferred to 250 mL pots containing the same substrate described above. Plants were maintained in the dark at 25 °C for 3 days and then moved to light conditions (200 µmol m ² s ¹; 14/10 h photoperiod). *A. tumefaciens* cultures were harvested and resuspended in infiltration medium (10 mM MgCl, 10 mM MES, and 200 µM acetosyringone) and adjusted to an OD of 0.6.

### Phytohormone treatments

Gibberellin (GA_3_) and PAC (Sigma-Aldrich) treatments were applied to plants by soil drenching, as previously described (Silva et al., 2019). GA_3_ treatments (10^-5^ M) of MT and SP5G^pen^ plants were performed from 1 to 10 days post-germination (DPG), while PAC treatments (10^-6^ M) were performed from 1 to 20 DPG. For both treatments, 25 mL of the solution was applied daily per pot. Germination was monitored individually to define ‘1 DPG’ (visible hypocotyl hook) (Figure 20). Ten-day-old treated seedlings were collected and subjected to RNA extraction as described above to evaluate the efficacy of the treatment.

To evaluate the impact of gibberellin on SP5G protein levels, the *SP5G::CFP* reporter was transiently expressed in *N. benthamiana* leaves via *Agrobacterium*-mediated infiltration. Plants were treated daily with mock or GA_3_ for three days by soil drenching and foliar spraying, using the same concentration applied in tomato treatments. At three days post-infiltration, leaf discs from each treatment were excised and placed in water in a 96-well black plate, with the abaxial side facing up. CFP fluorescence was measured using a Tecan Infinite 200 Pro plate reader (excitation: 433 nm; emission: 480 nm).

### Phylogenetic analysis

The SBP domains of proteins from *Arabidopsis* and tomato were aligned using ClustalW (http://www.ebi.ac.uk/Tools/clustalw). A maximum-likelihood phylogeny was then inferred in MEGA 6 (Kumar et al., 2016) with 1,000 bootstrap replicates, using default parameters.

### Generative AI Usage Statement

During the preparation of this work, the authors used the ChatGPT generative AI tool (OpenAI, GPT-5.3) to improve the clarity and accuracy of the language. After using this tool, the authors carefully reviewed and revised the content where necessary and take full responsibility for the final version of the manuscript.

### Statistical analysis

Statistical analysis was performed using GraphPad Prism 8 (GraphPad Software Inc., San Diego, CA, USA). Exact sample sizes (n) are indicated in each figure. Data were subjected to ANOVA, with means compared by the Student’s t-test or Tukey’s test as appropriate. When ANOVA assumptions were not met, non-parametric comparisons were made using the Mann–Whitney U or Dunn’s test.

## Accession numbers

*PHYB1*, Solyc01g059870; *TM3*, Solyc01g093965; *CNR*, Solyc02g077920; *SlCOL1*, Solyc02g089540; *SlCOL2*, Solyc02g089500; *SlCOL3*, Solyc02g089520; *SlGA20ox1*, Solyc03g006880; *DST*, Solyc03g006900; *SFT*, Solyc03g063100; *FUL2*, Solyc03g114830; *SlSBP6a*, *Solyc03g114850*; *FA*, Solyc03g118160; *ACTIN4*, Solyc04g011500; *BETA-TUBULIN 1*, Solyc04g081490; *ALPHA-TUBULIN 3*, Solyc04g077020; *SlSBP10*, Solyc05g015510; *SlSBP13*, Solyc05g015840; *SP5G*, Solyc05g053850; *SlGA20ox2*, Solyc06g035530; *FUL1*, Solyc06g069430; *SlGA2ox1*, Solyc07g056670; *SlSBP4b*, Solyc07g062980; *SlSBP3a*, Solyc10g009080; *SlSBP15*, Solyc10g078700; *PRO/DELLA*, Solyc11g011260, and *SlSBP6c*, Solyc12g038520

## Author contributions

M.H.V. and F.T.S.N. conceived, planned, and designed the experimental approach. M.H.V., G.S.B., F.G.P., R.R., A.C.F.F., R.G., Y.N.C.G., and C.C.F. conducted the experiments. K.P. and M.B. performed the EMSA assays. L.E.P. carried out the plant transformations. P.R. and F.B. provided the RNA-seq analysis. M.H.V., F.T.S.N., and F.V. directly supervised the experimental works. M.H.V. and F.T.S.N. wrote the manuscript, with contributions to revision by F.V., L.E.P.P., M.B., and M.C. All authors reviewed and approved the final version of the manuscript.

## Acknowledgments

We thank the members of Dr Nogueira and Dr Valverde laboratories for helpful discussions. Cassia Regina Figueiredo and Isabel María Jiménez Beníte for excellent technical support. We thank the FAPESP Multiuser Equipment Program (grant no. 19/27255-5) for supporting the plate reader used in this study.

## Data availability

The raw sequence data generated in this work have been deposited in the NCBI Gene Expression Omnibus (GEO) under the number GSE310414. Correspondence and requests for materials should be addressed to F.T.S.N and M.H.V.

