## Supplemental Figures for "*SELF-PRUNING 5G* interplays with age and gibberellin pathways downstream of *SlCOL1* to orchestrate photoperiodic tomato flowering"

Running title: COL1–SP5G links photoperiod, age, and GA Pathways

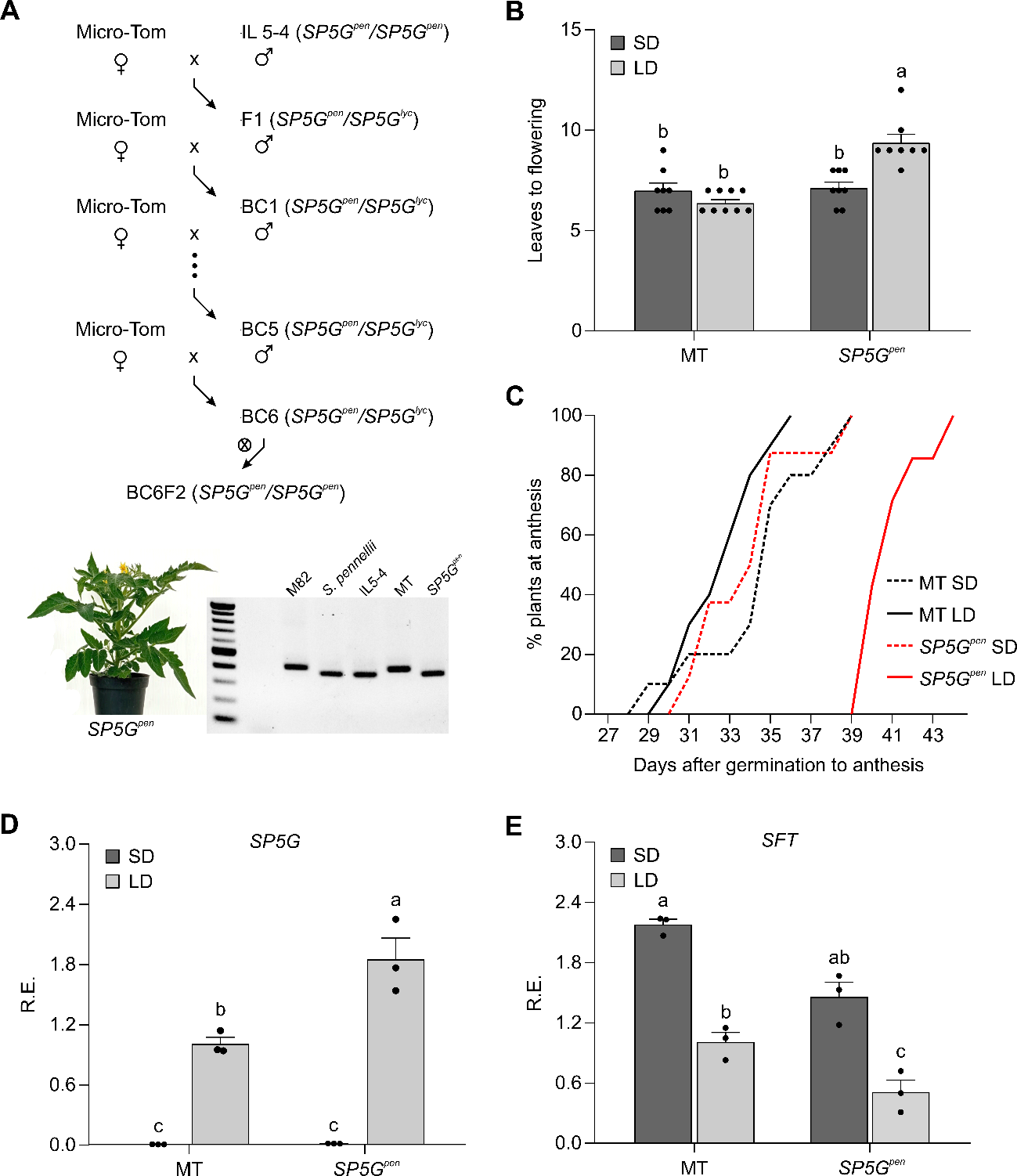

**Figure S1. The functional *SP5G* allele from the wild species *S. pennellii* represses flowering in cultivated tomato under LD. (A)** Top panel: Crossing scheme to obtain the *SP5G^pen^* genotype in the tomato cv. Micro-Tom (MT) background. IL5-4 represents an introgression line generated by Eshed and Zamir (1995) in the tomato cv. M82 background. Bottom panel: *left*, representative image of an *SP5G^pen^* plant (at 40 days post-germination); *right*, agarose gel electrophoresis for the molecular differentiation of *SP5G* alleles from *S. lycopersicum* (size = 411 bp) and *S. pennellii* (size = 378 bp). **(B)** Leaves to flowering of MT and *SP5G^pen^* genotypes under short day (SD, 10/14h, dark grey bars) and long day (LD, 14/10h, light grey bars) conditions (mean ± SEM; Tukey’s test, *p<0.001*; n=8 plants). Each dot represents an individual data. **(C)** Percentage of plants at anthesis after germination in SD (dashed line) and LD (solid line) conditions for MT (black line) and *SP5G^pen^* (red line) plants (n=8 plants). **(D and E)** Relative expression (R.E.) of *SP5G* (**D**) and *SFT* (**E**) genes in MT and *SP5G^pen^* genotypes under SD (dark grey bars) and LD (light grey bars) conditions (ZT=4; mean ± SEM; Distinct letters indicate significant differences according to Tukey’s test, *p<0.05*; n=3 samples). ZT=4 was chosen as the time point for all RT-qPCR analyses because it corresponds to the peak of *SP5G* expression (Soyk et al., 2017; Zhang et al., 2018). Each dot represents an individual data.

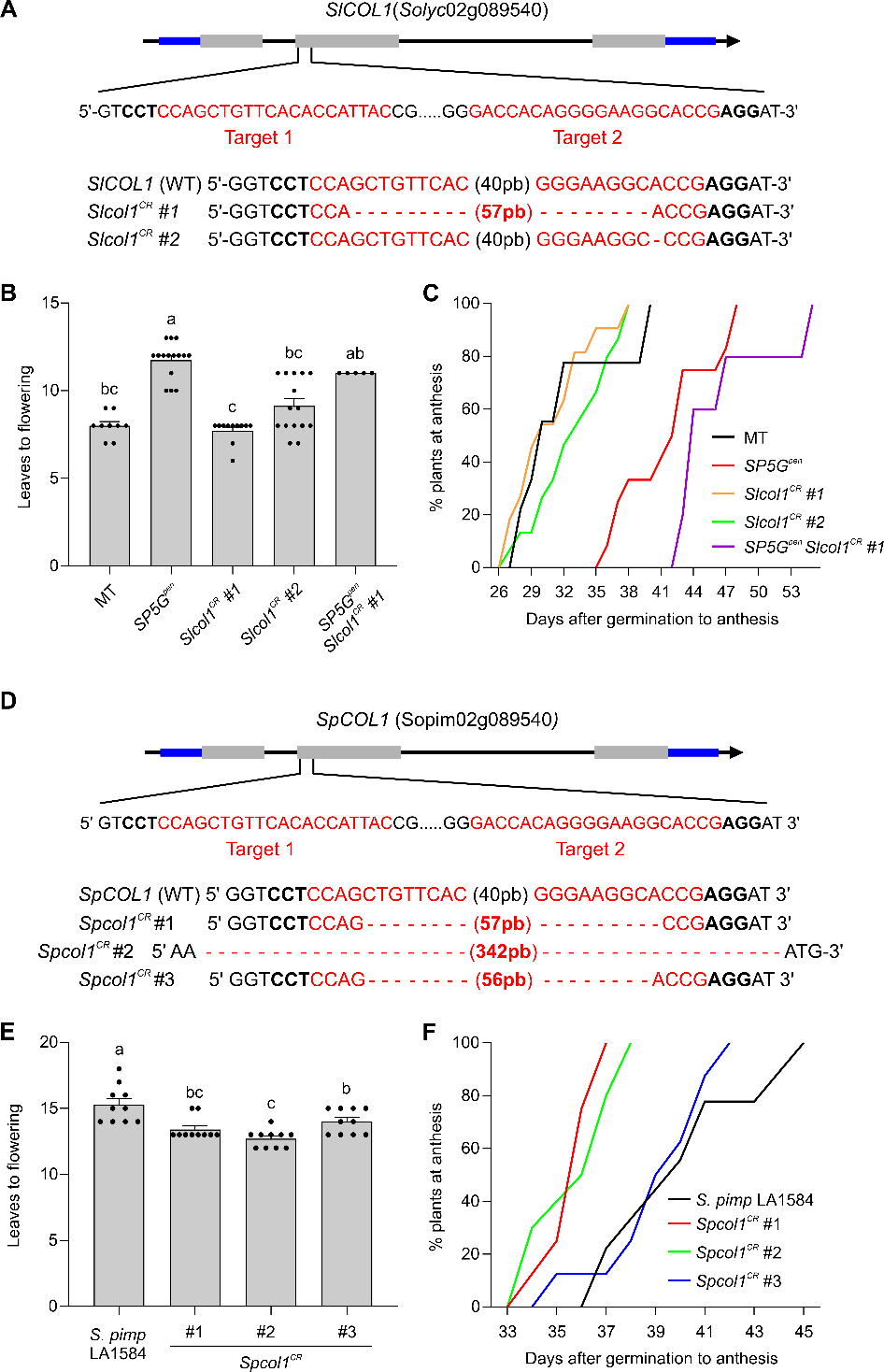

**Figure S2. Molecular and phenotypic characterization of *COL1* mutants in MT and *S. pimpinellifolium*** **generated by CRISPR-Cas9 technology. (A)** Top panel: *SlCOL1* was targeted with two single-guide RNAs (highlighted in red) for CRISPR/Cas9-induced gene editing. Bold font indicates the Protospacer-Adjacent Motif (PAM) sequences. Bottom panel: DNA sequencing shows the alleles generated by the CRISPR-Cas9 technology. *Slcol1^CR #1^* allele was characterized by a 57 bp deletion, while *Slcol1^CR #2^* allele showed a single adenine deletion at position 388. **(B and C)** Leaves to flowering (**B**) and percentage of plants at anthesis after germination (**C**) in LD (14/10h) conditions for MT, *SP5G^pen^*, *Slcol1^CR^ #1*, *Slcol1^CR^ #2*, and the double *SP5G^pen^* *Slcol1^CR^ #1* plants (mean ± SEM; Dunn’s test, p<0.01; n = 5-15 plants). **(D)** Top panel: The *COL1* from *S. pimpinellifolium* was targeted with the same two single-guide RNAs (highlighted in red) used to generate *Slcol1^CR^* mutants in tomato. Bold font indicates the Protospacer-Adjacent Motif (PAM) sequences. Bottom panel: DNA sequencing shows the alleles generated by the CRISPR/CAS9 approach. Red numbers in parentheses indicate the sizes of the deletions in the CRISPR-Cas9 lines. **(E and F)** Leaves to flowering (**E**) and percentage of plants at anthesis after germination (**F**) in LD (14/10h) conditions for *S. pimpinellifolium* (accession LA1584) and *Spcol1 CRISPR* lines (*Spcol1^CR^*) (mean ± SEM; Tukey’s test, p<0.01; n = 8-10 plants). Distinct letters indicate significant differences according to the applied test. In (**B**) and (**E**), each dot represents an individual data.

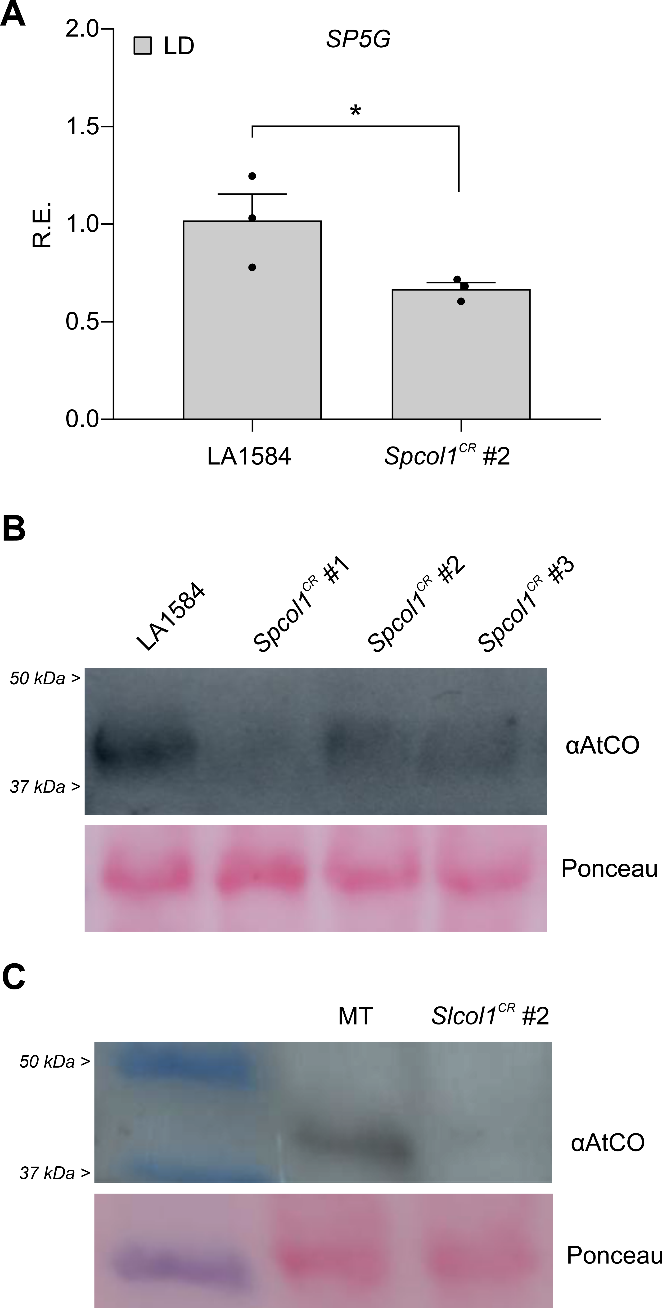

**Figure S3. *Spcol1^CR^* mutants showed reduced levels of *SP5G* transcripts and SpCOL1 protein.** **(A)** Relative expression (R.E.) of the *SP5G* gene in *S.* *pimpinellifolium* (LA1584) and *Spcol1^CR^* (allele #2) seedlings at 10 DPG in LD (ZT = 4; mean ± SEM; Student’s *t*-test, *p<0.05*; n = 3 samples). Each dot represents an individual data. **(B)** Immunoblot analysis of SpCOL1 protein accumulation in seedlings of *S. pimpinellifolium* (LA1584) and *Spcol^CR^* mutants (allele #1, #2, and #3) at 10 DPG. **(C)** Immunoblot analysis of SlCOL1 protein accumulation in MT and *Slcol1^CR^* (allele *#2*) seedlings at 10 DPG. Samples were harvested at ZT4. Membranes were incubated with αAtCO. Ponceau staining was used as a loading control.

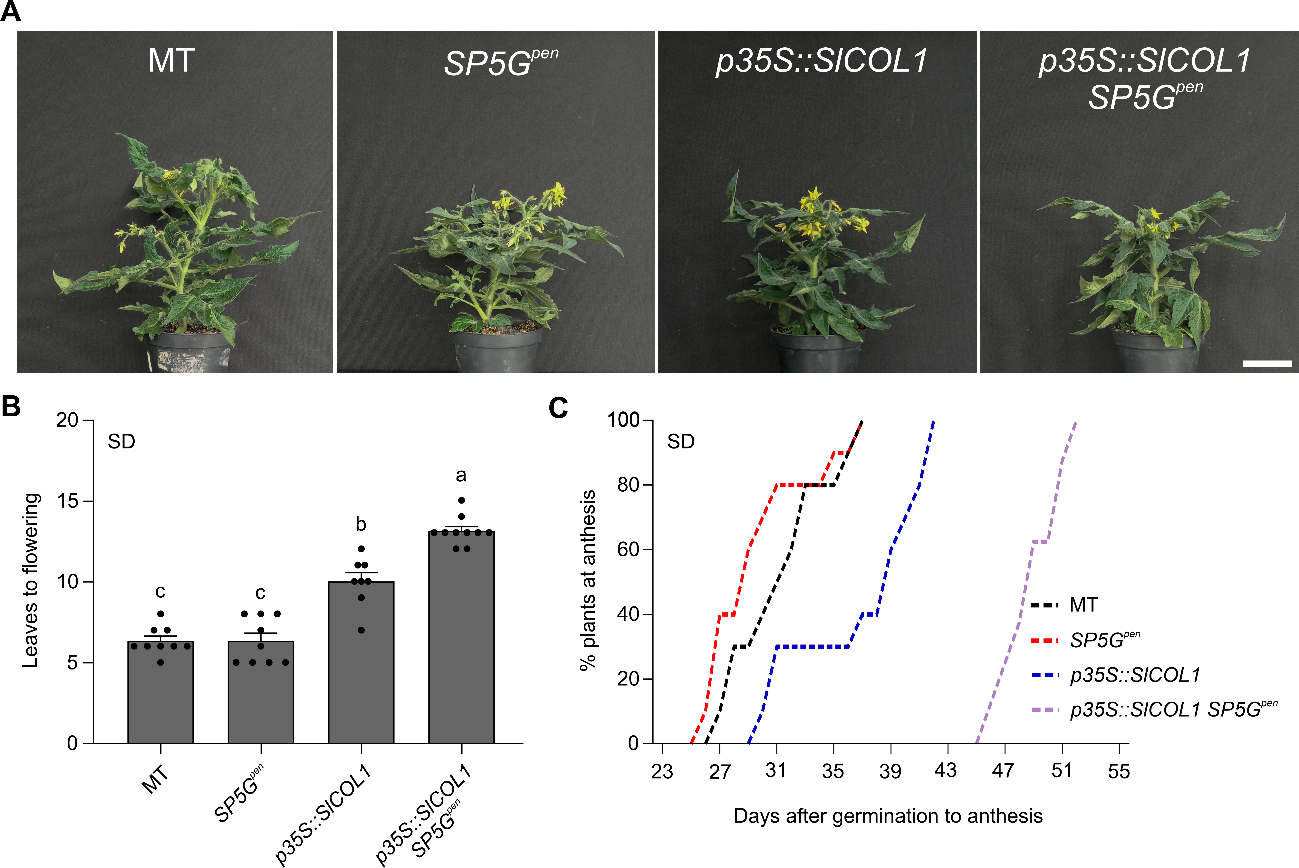

**Figure S4. High levels of *SlCOL1* repress tomato flowering in SD, especially in plants harboring the *SP5G^pen^* allele. (A)** Representative pictures of 50-day-old plants of MT, *SP5G^pen^*, *p35S::SlCOL1*, and *p35S::SlCOL1 SP5G^pen^* cultivated in SD (10h/14h). Bar = 5cm. **(B)** Leaves to flowering of MT, *SP5G^pen^*, *p35S::SlCOL1*, and *p35S::SlCOL1 SP5G^pen^* cultivated in SD. Data are expressed as mean ± SEM (Tukey’s test, *p<0.001*; n = 8-10 plants). Each dot represents an individual data. **(C)** Percentage of plants at anthesis after germination in SD conditions for MT (black line), *SP5G^pen^* (red line), *p35S::SlCOL1* (blue line), and *p35S::SlCOL1 SP5G^pen^* (purple line) plants (n = 8-10 plants).

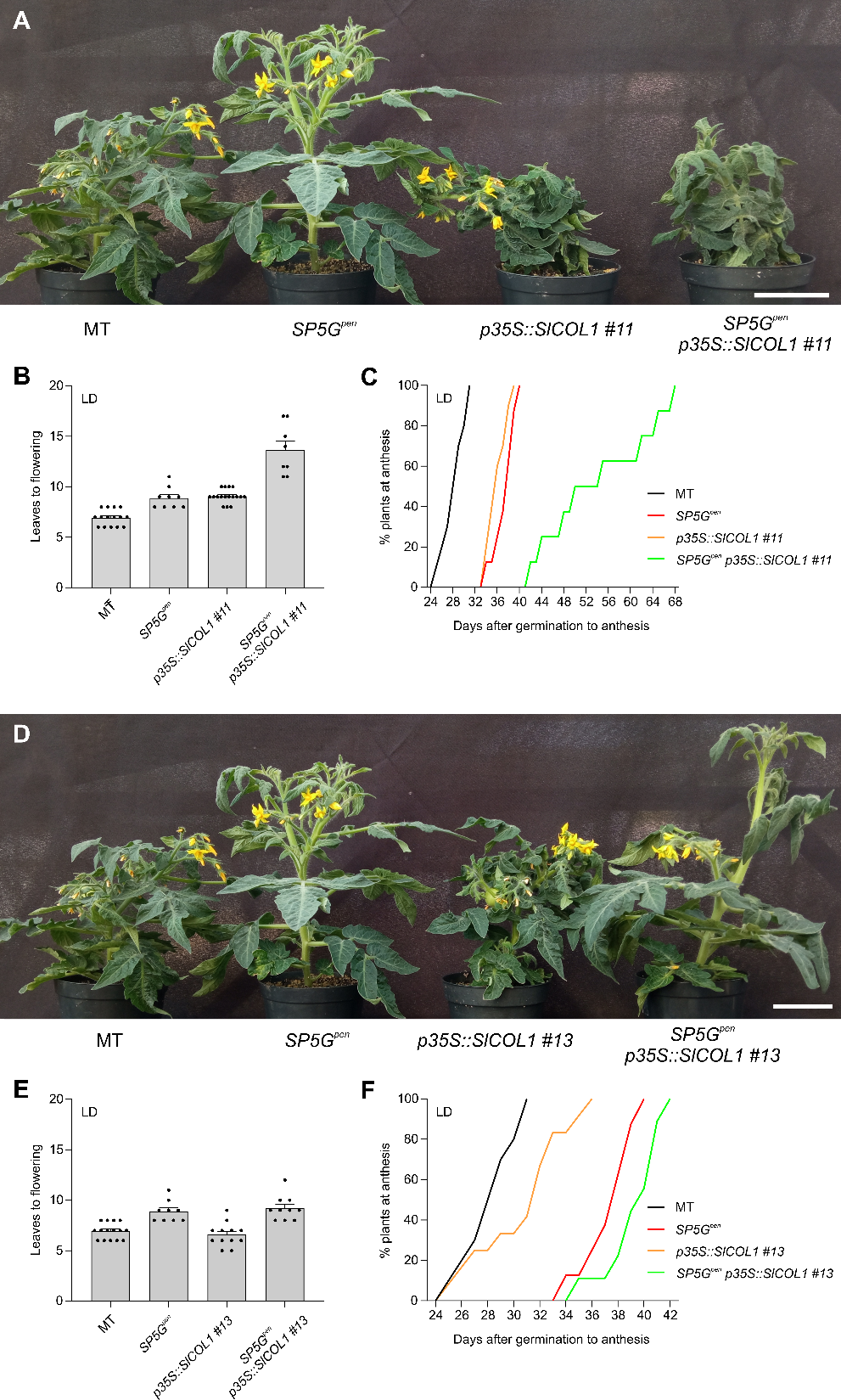

**Figure S5. Delaying tomato flowering in an *SP5G*-dependent manner requires high levels of SlCOL1 protein. (A)** Representative pictures of 45-day-old plants of MT, *SP5G^pen^*, *p35S::SlCOL1* line #11, and the double *p35S::SlCOL1* line #11 x *SP5G^pen^* grown in LD (14h/10h). Bar = 5 cm. **(B and C)** Leaves to flowering (**B**) and percentage of plants at anthesis after germination (**C**) in LD conditions for MT, *SP5G^pen^*, *p35S::SlCOL1* line #11, and the double *p35S::SlCOL1* line #11 x *SP5G^pen^* plants. The data are expressed as mean ± SEM (Tukey’s test, p<0.05; n = 8-17 plants). **(D)** Representative pictures of 45-day-old plants of MT, *SP5G^pen^*, *p35S::SlCOL1* line #13, and the double *p35S::SlCOL1* line #13 x *SP5G^pen^* grown in LD (14h/10h). Bar = 5cm. **(E and F)** Leaves to flowering (**E**) and percentage of plants at anthesis after germination (**F**) in LD conditions for MT, *SP5G^pen^*, *p35S::SlCOL1* line #13, and the double *p35S::SlCOL1* line #13 x *SP5G^pen^* plants. The data are expressed as mean ± SEM (Tukey’s test, p<0.05; n = 9-15 plants). In (**B**) and (**E**), each dot represents an individual data.

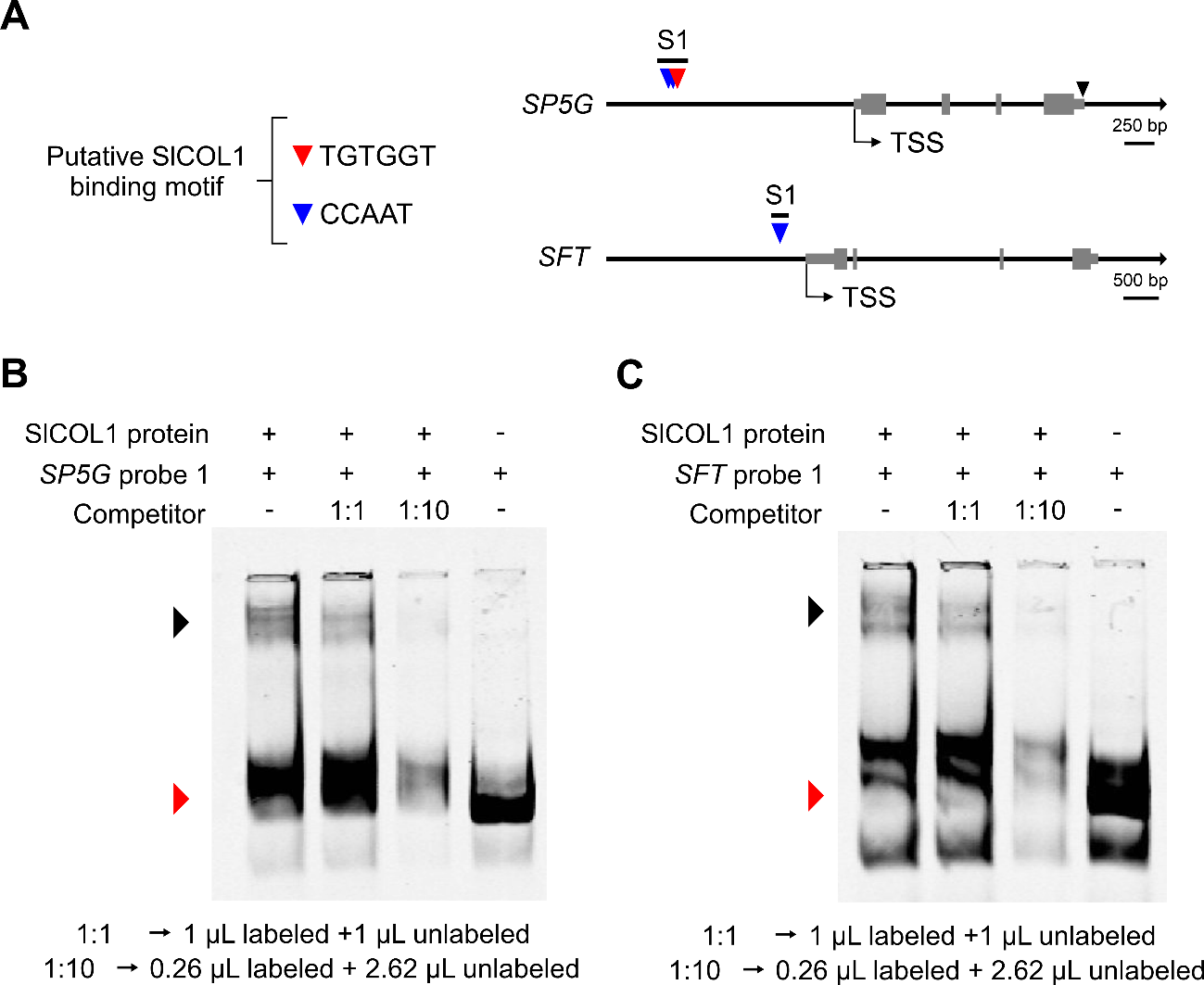

**Figure S6. SlCOL1 binding to the *SP5G* and *SFT* promoter fragments. (A)** Schematic representations of the *SP5G* (top) and *SFT* (bottom) loci showing the putative SlCOL1-binding motifs. Grey boxes represent exons. TSS indicates the transcription start site. Red and blue triangles indicate putative SlCOL1 binding motifs, TGTGGT and CCAAT, respectively. The black triangle marks the position of the 52-bp deletion. **(B and C)** EMSA assays testing the ability of SlCOL1 to bind to regulatory regions of *SP5G* (**B**) and *SFT* (**C**) genes. The black arrowhead indicates the protein-probe complex, and the red arrowhead indicates the free probe.

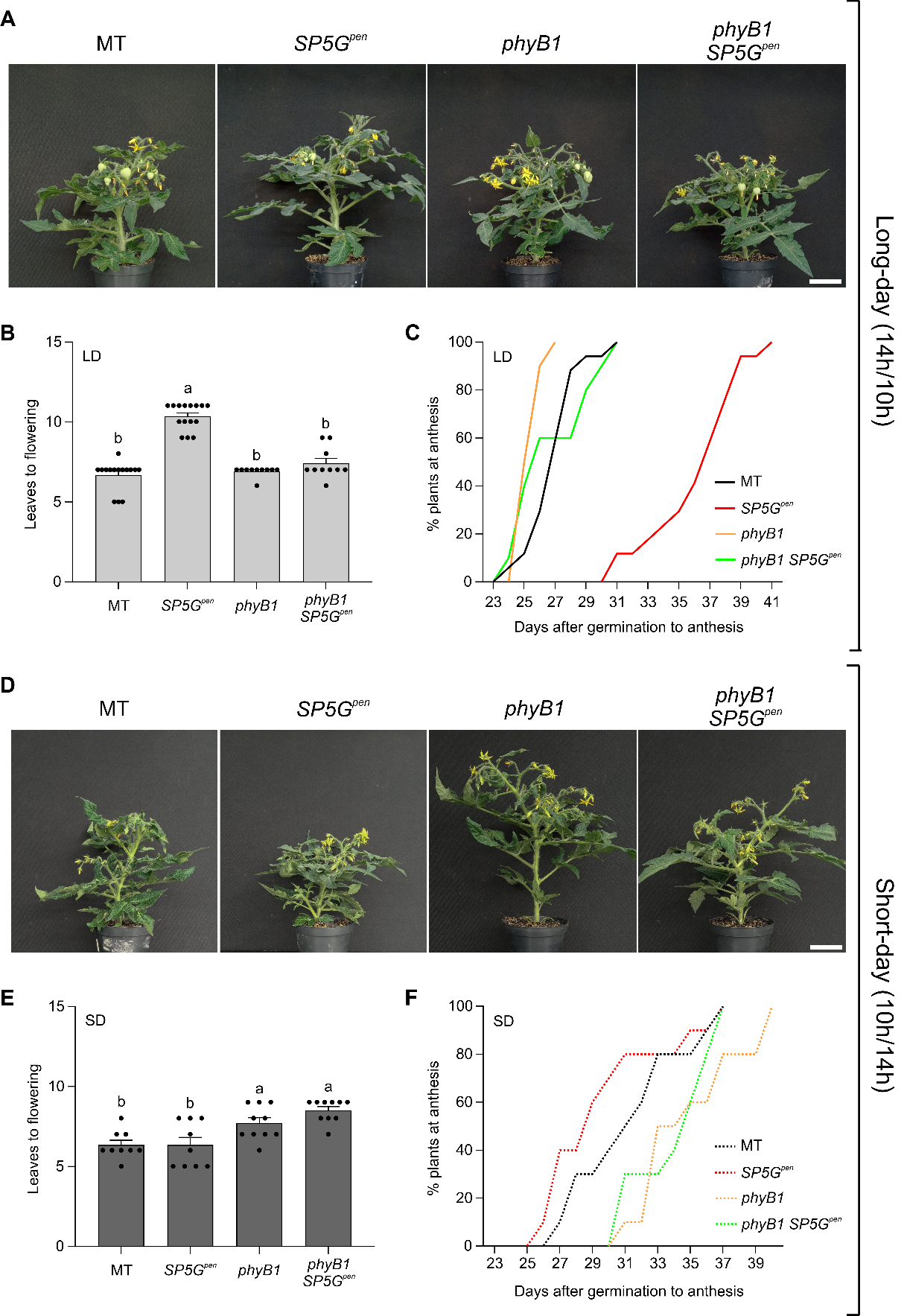

**Figure S7. *phyB1* mutation rescues the late flowering phenotype of *SP5G^pen^* plants in LD. (A and D)** Representative pictures of MT, *SP5G^pen^*, *phyB1*, and *phyB1 SP5G^pen^* cultivated in LD (**A**, 14h/10h) and SD (**D**, 10h/14h). Bar = 5cm. Plants were 60- and 50-day-old at the time of imaging under LD and SD conditions, respectively. **(B and E)** Leaves to flowering of MT, *SP5G^pen^*, *phyB1*, and *phyB1 SP5G^pen^* in LD (**B**) and SD (**E**). Data are expressed as mean ± SEM (Tukey’s test, p<0.001; n = 10-15 plants). **(C and F)** Percentage of plants at anthesis after germination for MT (black line), *SP5G^pen^* (red line), *phyB1* (orange line), and *phyB1 SP5G^pen^* (green line) plants cultivated in LD (**C**) and SD (**F**) (n=10-15 plants). Solid and dashed lines represent plants grown in LD and SD conditions, respectively. Distinct letters indicate significant differences according to the applied test. In (**B**) and (**E**), each dot represents an individual data. Controls shown in Figure S7A–7C are the same as those used in Figure 2A–2C, whereas the controls in Figure S7D–7E correspond to those presented in Figure S4A–4C.

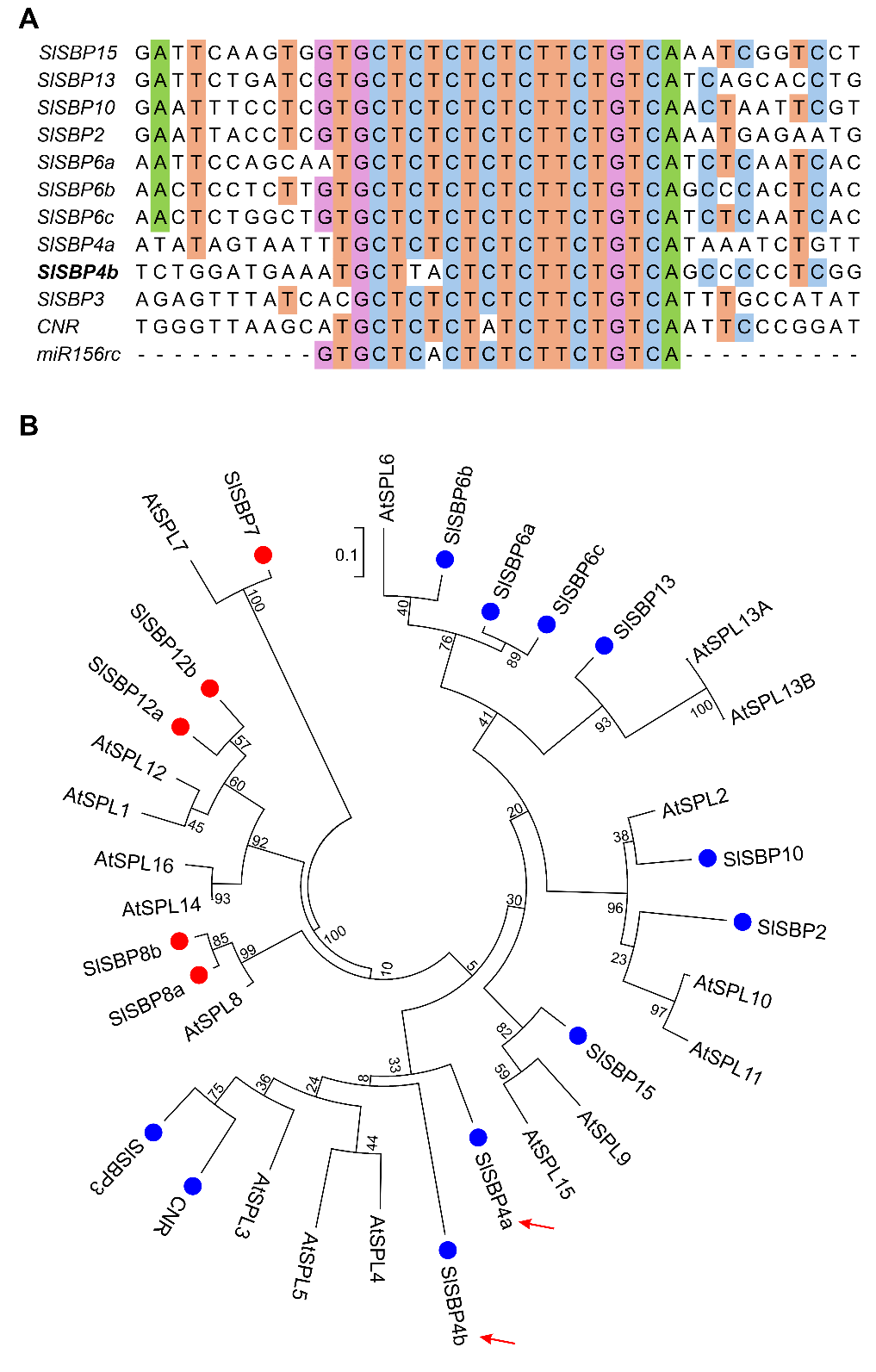

**Figure S8. miR156-targeted *SlSBP4b* is a new member of the tomato SPL/SBP transcription factor family. (A)** Sequence alignment of miR156 complementary sites in *SlSBP* genes. miR156rc indicates the reverse complement sequences of the mature miR156, which was used in the alignment for comparison. **(B)** Phylogenetic analysis of SBP-domain proteins in *Arabidopsis* (AtSPLs) and tomato (SlSBPs). Conserved SBP domains were aligned, and a maximum‐likelihood tree was inferred in MEGA6 using 1,000 bootstrap replicates. Branch lengths (scale bar = 0.1) and bootstrap support values are shown. Blue dots indicate *SlSBP* genes targeted by miR156, while red dots represent non-targeted SBPs. Red arrows indicate the previously annotated *SBP4*, here renamed *SlSBP4a*, and its newly identified paralog, *SBP4b*.

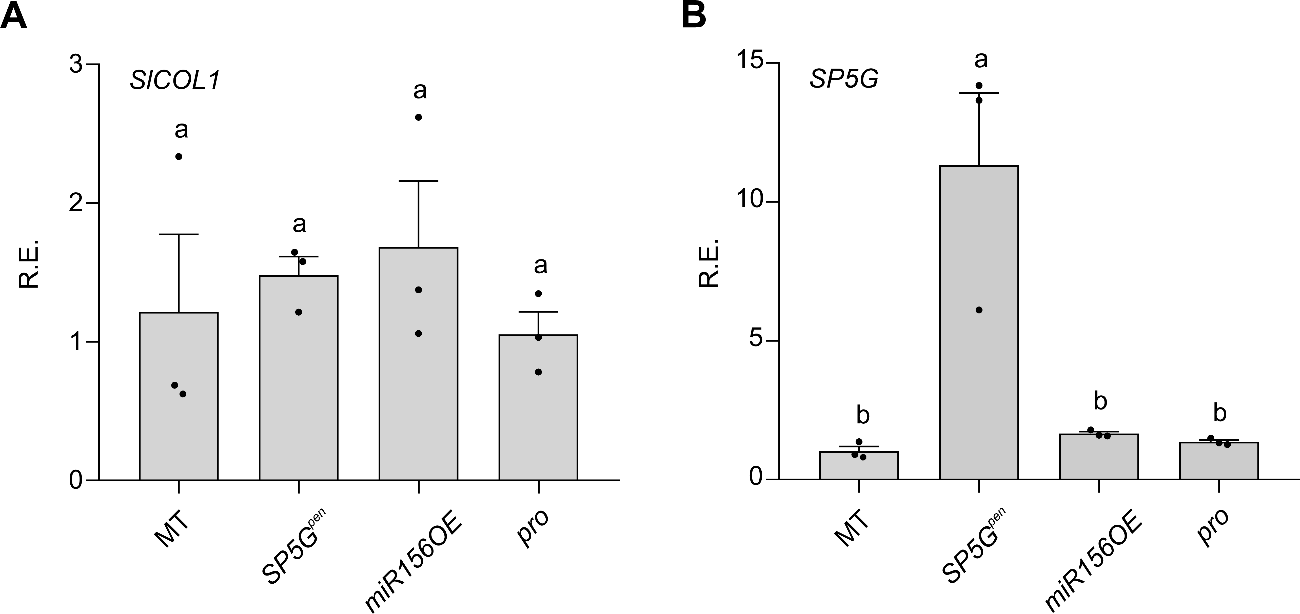

**Figure S9. *SlCOL1* and *SP5G* expression in genotypes with modified photoperiod, age, and PRO/DELLA pathways. (A and B)** Relative expression of *SlCOL1* (**A**) and *SP5G* (**B**) genes in MT, *SP5G^pen^*, *miR156OE*, and *procera* (*pro*) genotypes under LD conditions (ZT = 4; mean ± SEM; n = 3 samples). Distinct letters indicate significant differences according to Tukey’s test, *p<0.001*. Each dot represents an individual data.

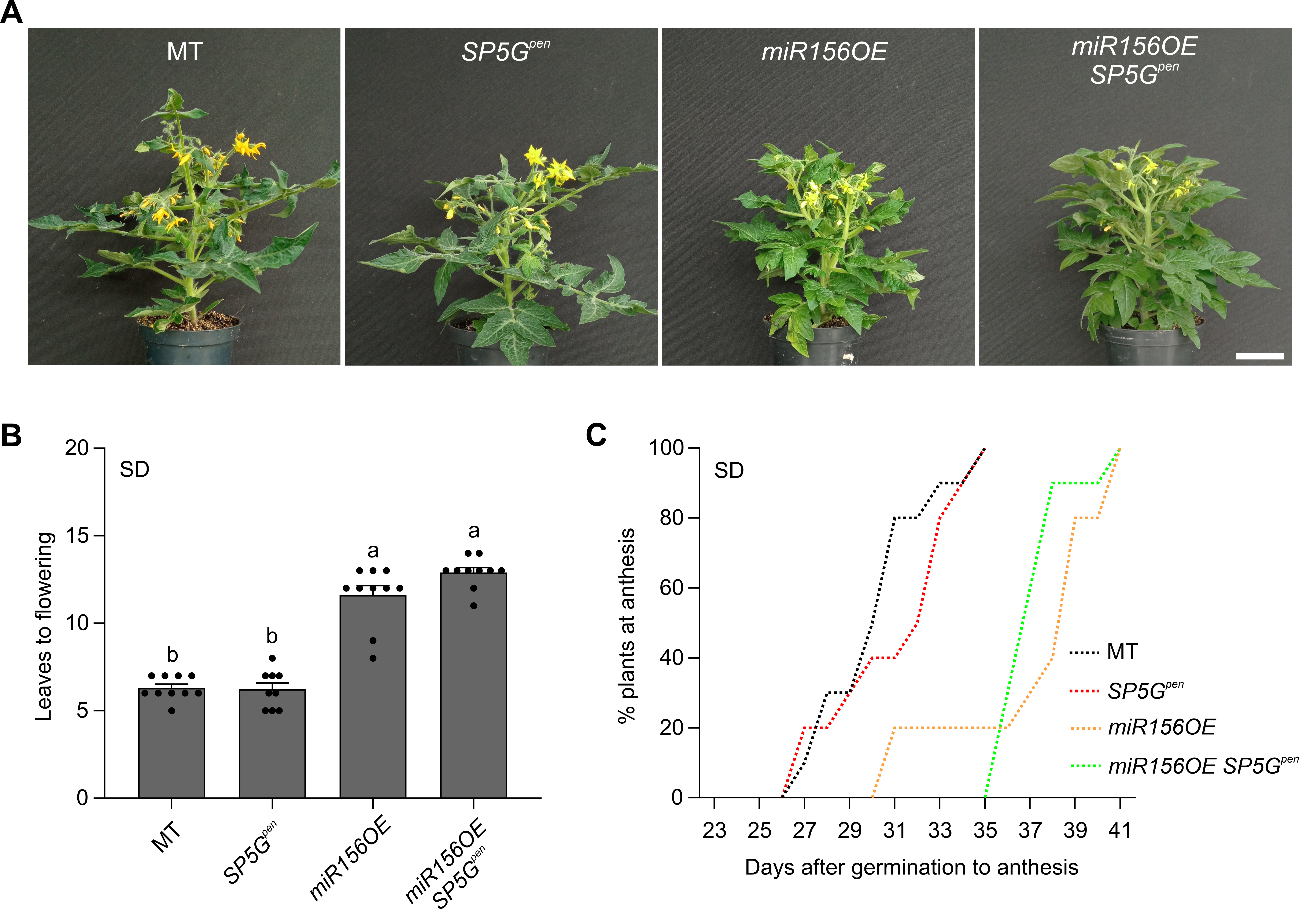

**Figure S10. No synergistic effect on flowering time is observed in *miR156OE* *SP5G^pen^* plants grown under SD conditions. (A)** Representative pictures of 50-day-old plants of MT, *SP5G^pen^*, *miR156OE*, and *miR156OE SP5G^pen^* cultivated in SD (10h/14h). Bar = 5cm. **(B and C)** Leaves to flowering (**B**) and percentage of plants at anthesis after germination (**C**) in SD conditions for MT, *SP5G^pen^*, *miR156OE*, and *miR156OE SP5G^pen^*. Data are expressed as mean ± SEM (n = 9-10 plants). Distinct letters indicate significant differences according to Tukey’s test (*p<0.001*). Each dot represents individual data.

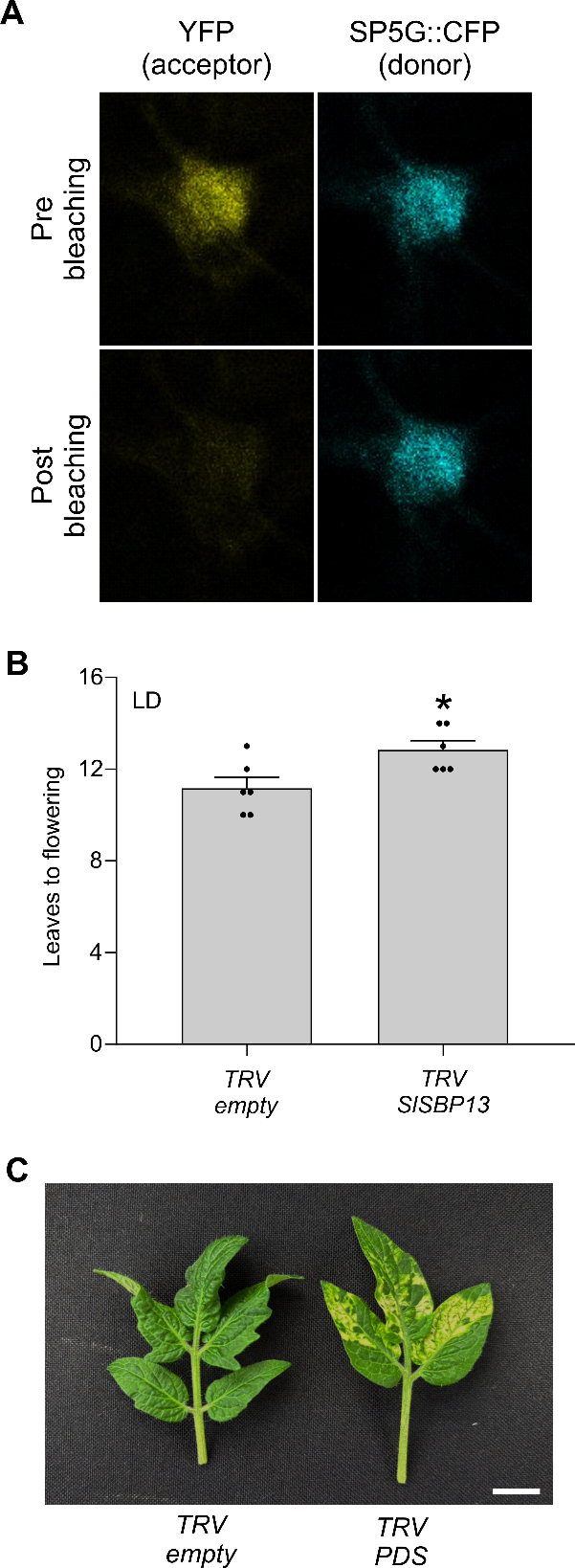

**Figure S11. VIGS-mediated silencing of *SlSBP13* delays flowering in tomato plants carrying the *SP5G^pen^* allele. (A)** Detection of FRET by acceptor photobleaching. Representative images of YFP and SP5G::CFP signals before and after photobleaching. **(B)** Leaves to flowering in *SP5G^pen^* plants infected with TRV-*SlSPB13* under LD conditions. Data are expressed as mean ± SEM (Student’s *t*-test, *p<0.05*; n = 6 plants). Each dot represents individual data. **(C)** Representative image showing leaf bleaching in *SP5G^pen^* plants infected with TRV-*SlPDS* as a positive control. Bar = 1 cm.

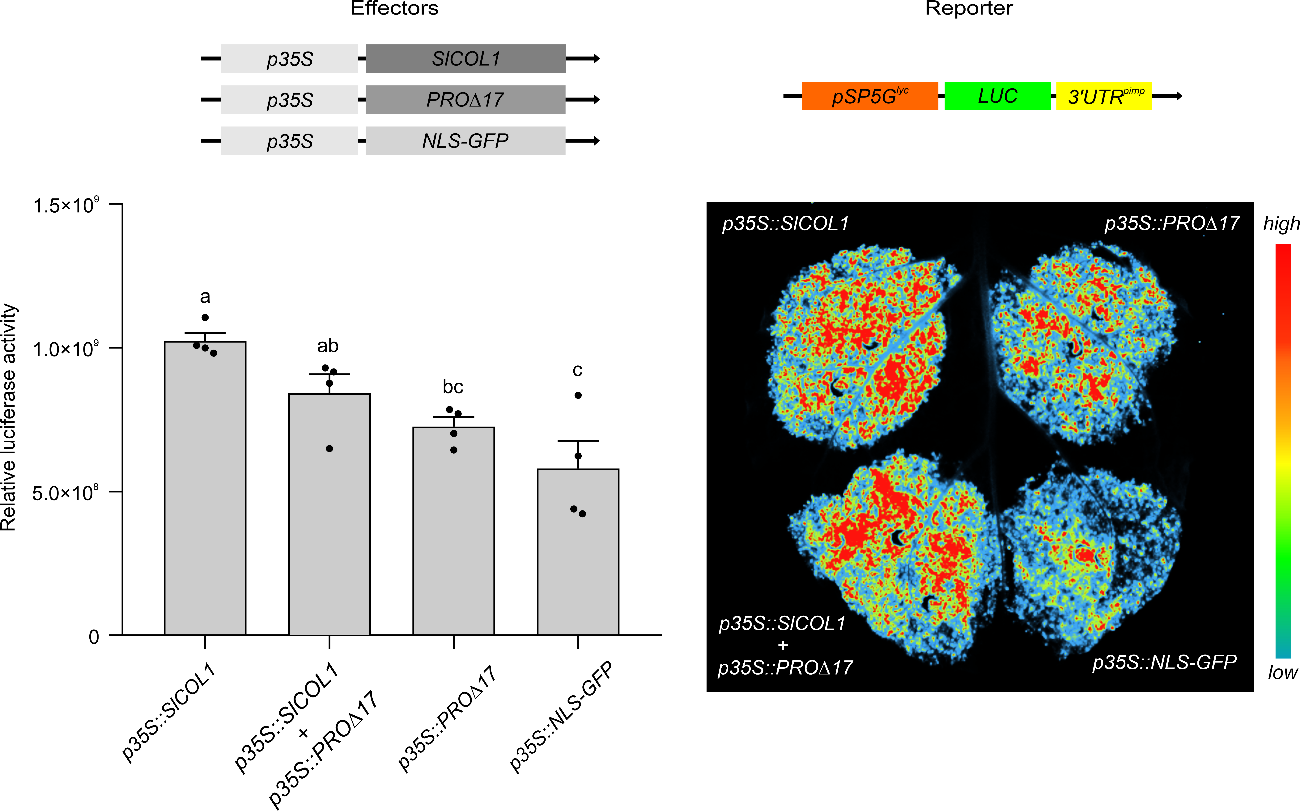

**Figure S12. Stable PROCERA/DELLA does not alter SlCOL1-dependent activation of *SP5G*.** Transient activation of *pSP5G^lyc^::LUC::3’UTR^pimp^* reporter constructs in *N. benthamiana* leaves. Four leaves were co-infiltrated with *p35S::SlCOL1*, *p35S::PRO∆17*, and/or *p35S::NLS-GFP* effector constructs. PRO∆17 is a protein with a deletion of the N-terminal 17 amino acids in the DELLA domain, resulting in its stabilization in response to GA. *Top:* Schematic representation of effector and reporter constructs used in this assay. *Bottom left:* Quantification of relative luciferase activity using NEWTON 7.0 imaging software. Values are mean ±SEM (n = 3). Distinct letters indicate significant differences according to Tukey’s test (*p<0.05*). *Bottom right:* Representative bioluminescence images captured using the NEWTON 7.0 CCD imaging system. Each dot represents an individual data.

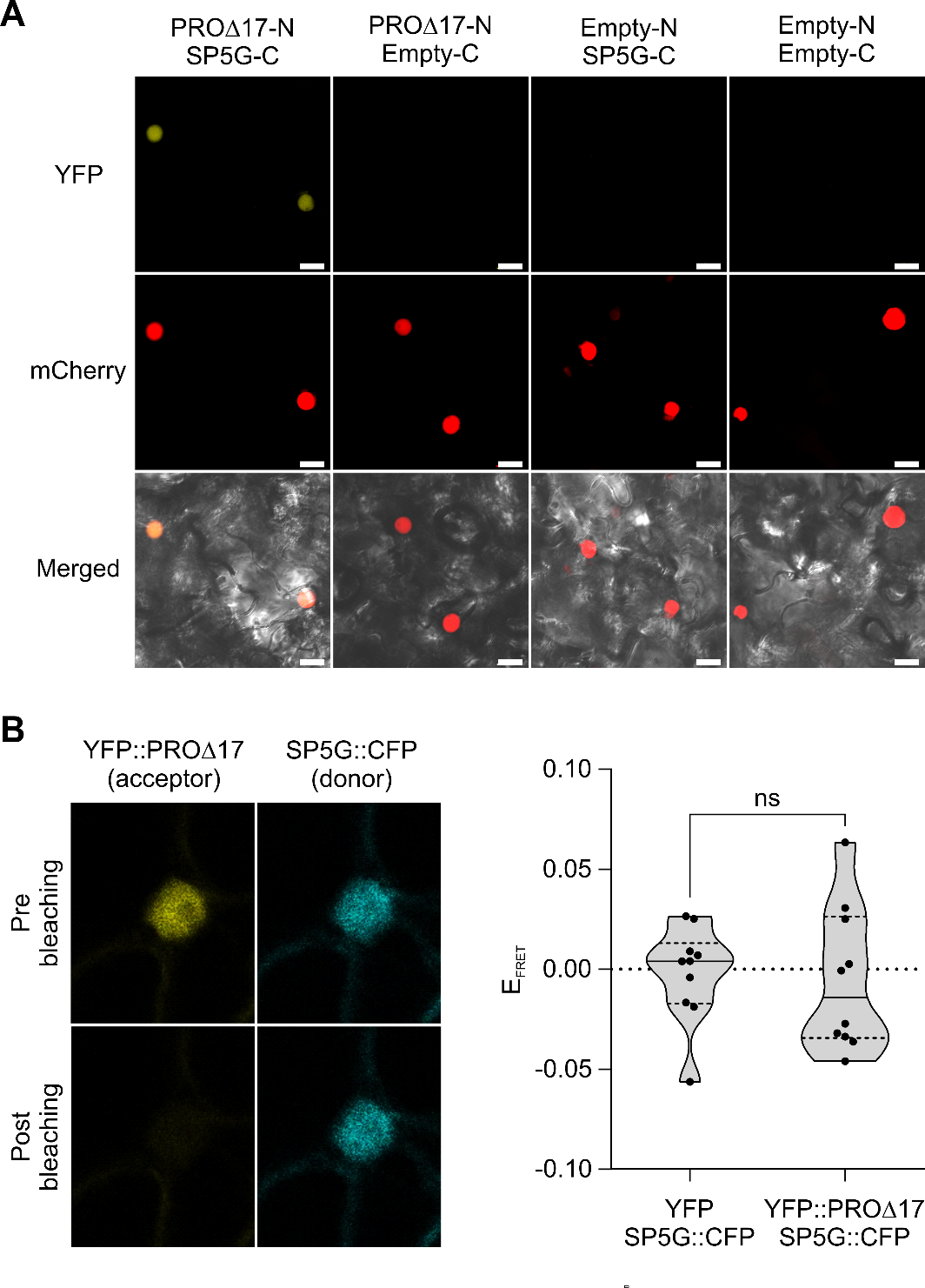

**Figure S13. SP5G and PRO physical interaction by BiFC. (A)** BiFC experiments using *N. benthamiana* leaves infiltrated with agrobacteria. N and C represent the N-terminal and C-terminal fragments of the YFP protein, respectively. Empty indicates infiltration with an empty vector. Combinations of PRO∆17-N with Empty-C, Empty-N with SP5G-C, and Empty-N with Empty-C were used as negative controls. *AtWWP1::mCherry* was used as a nuclear marker. Bright field and merged images are also shown. Scale bars = 20 μm. **(B)** FRET acceptor photobleaching measurements between SP5G and PRO∆17. FRET efficiency (E_FRET_) was calculated as described in Fig. 5f. Values are mean ±SEM (Student’s *t*-test, ns = not significant, n = 10 cells). The same negative control (YFP co-infiltrated with *SP5G::CFP*) was used for both interaction assays (*SP5G* with *rSBP13* and *SP5G* with PRO∆17). Representative images of YFP::PRO∆17 and SP5G::CFP signals before and after photobleaching are shown (left). Images of the negative control are presented in Figure S11. Each dot represents an individual data.

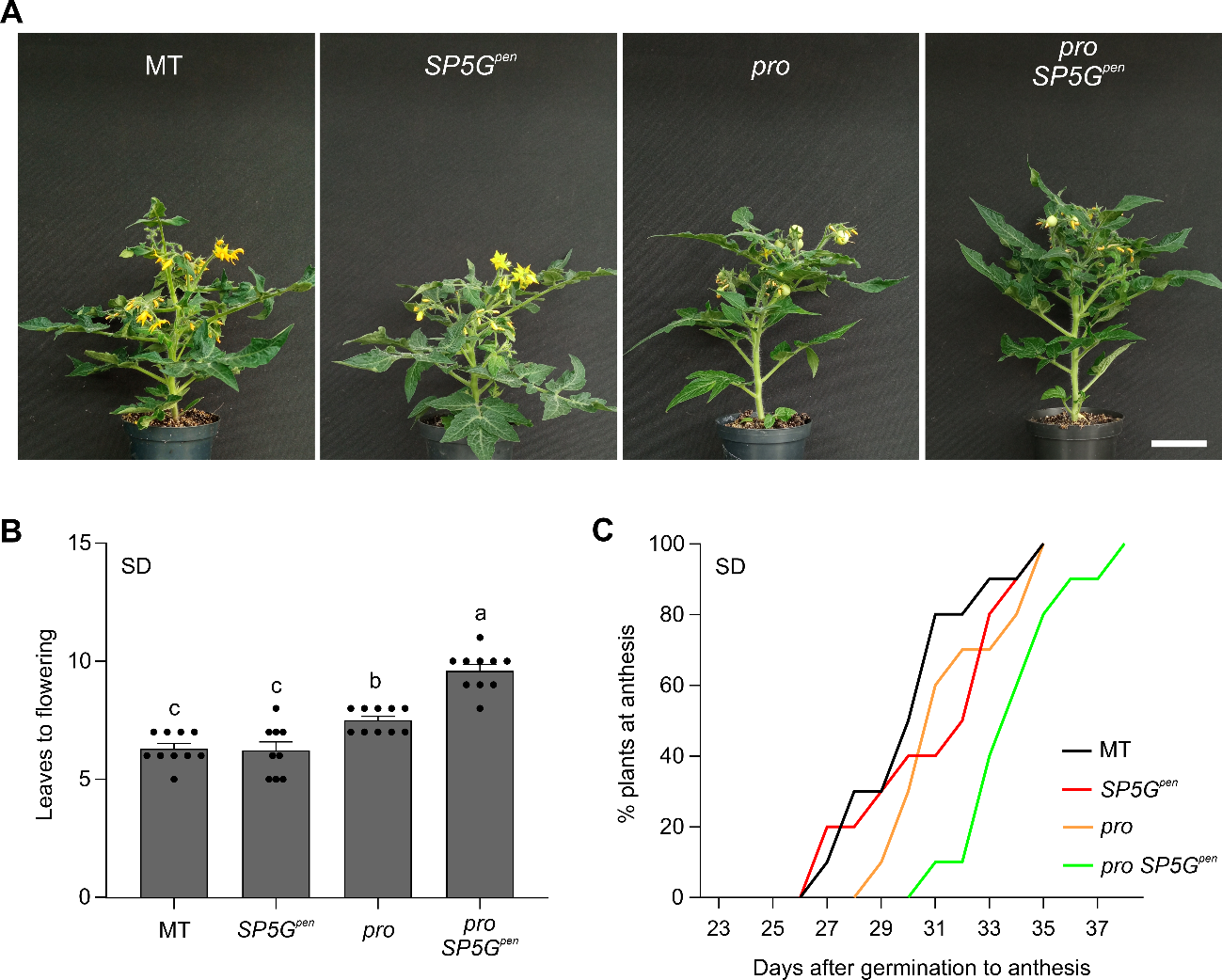

**Figure S14. *pro SP5G^pen^* plants exhibited a synergistic delay in flowering under SD conditions. (A)** Representative pictures of 50-day-old plants of MT, *SP5G^pen^*, *pro*, and *pro SP5G^pen^* cultivated in SD (10h/14h). Bar = 5cm. **(B and C)** Leaves to flowering (**B**) and percentage of plants at anthesis after germination (**C**) in SD conditions for MT, *SP5G^pen^*, *pro*, and *pro SP5G^pen^*. Data are expressed as mean ± SEM (n = 9-10 plants). Distinct letters indicate significant differences according to Tukey’s test (*p<0.001*). Each dot represents an individual data. Controls shown in Figure S14A–14C are the same as those used in Figure 10A–10C.

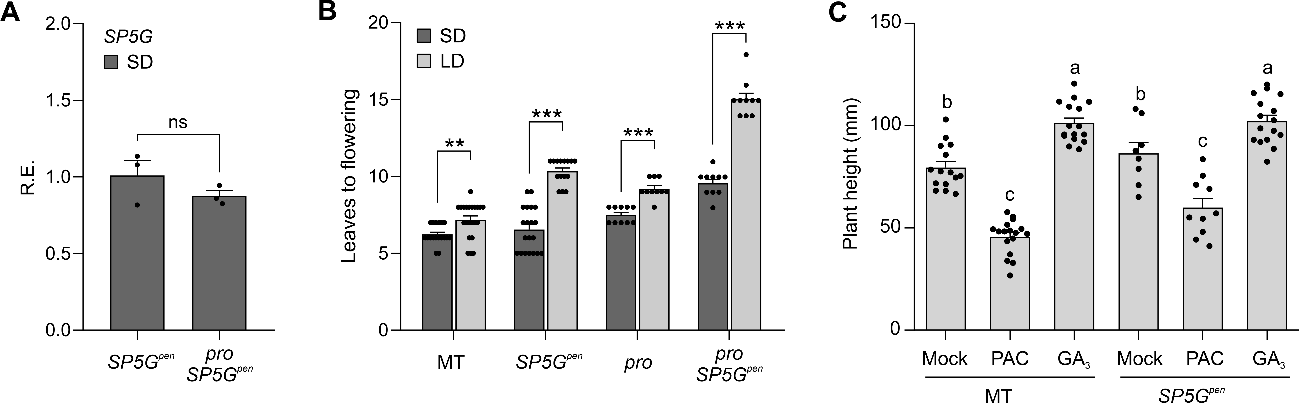

**Figure S15. *SP5G*-dependent photoperiodic flowering is enhanced in the absence of functional PRO/DELLA. (A)** Relative expression of *SP5G* gene in *SP5G^pen^* and *pro SP5G^pen^* genotypes under SD conditions (ZT=4; mean ± SEM; Student’s t test, *p<0.05*; ns = not significant; n=3 samples). **(B)** Leaves to flowering of MT, *SP5G^pen^*, *pro*, and *pro SP5G^pen^* genotypes in SD (10/14h, dark grey bars) and LD (14/10h, light grey bars). The data are expressed as ± SEM (Student’s t test, n=10-22 plants). Two (**) and three (***) asterisks indicate *p<0.01* and *p<0.001*, respectively. **(C)** Height to first inflorescence of MT and *SP5G^pen^* plants treated with mock, paclobutrazol (PAC), and gibberellin (GA_3_) in LD (Tukey’s test, *p<0.05*; n = 7-17). Distinct letters indicate significant differences according to the applied test. Each dot represents an individual data.

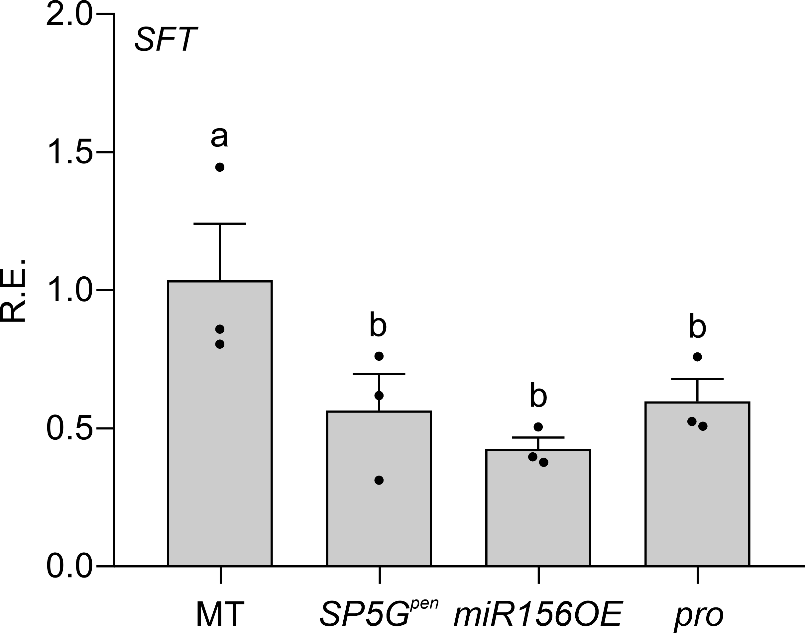

**Figure S16. Relative expression of *SFT* in MT, *SP5G^pen^*, *miR156OE*, and *pro* mature leaves in LD.** Distinct letters indicate significant differences according to Tukey’s test (ZT = 4; mean ± SEM; n = 3 samples, *p<0.05*). Each dot represents an individual data.

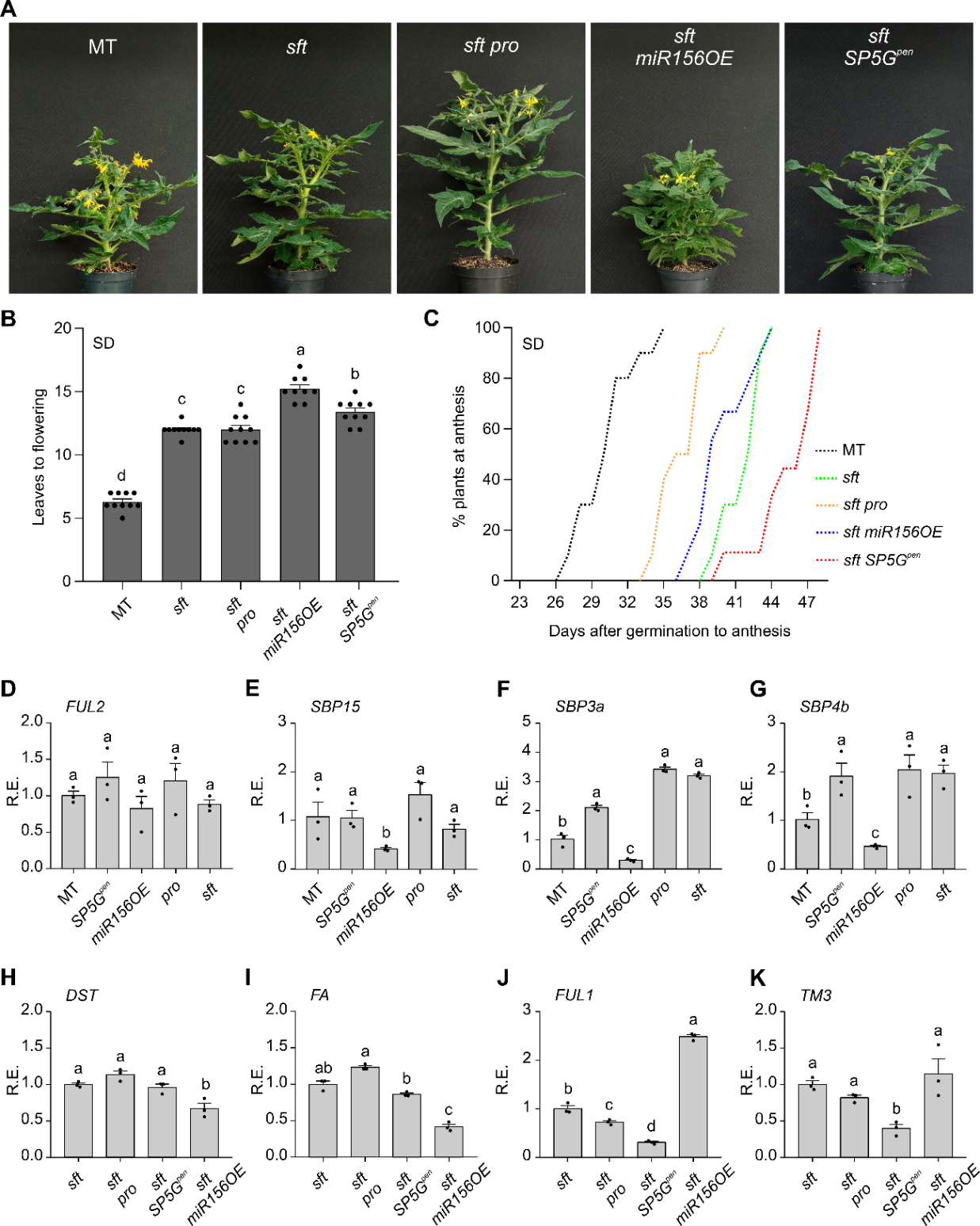

**Figure S17: *pro* and *sft pro* showed similar flowering time in SD. (A)** Representative pictures of 50-day-old plants of MT, *sft*, *sft pro*, *sft miR156OE*, and *sft SP5G^pen^* cultivated in SD (10h/14h). Bar = 5cm. **(B and C)** Leaves to flowering (**B**) and percentage of plants at anthesis after germination (**C**) in SD conditions for MT, *sft*, *sft pro*, *sft miR156OE*, and *sft SP5G^pen^* plants. Data are expressed as mean ± SEM (n = 9-10 plants). **(D-G)** Relative expression of *FUL2* (**D**), *SBP15* (**E**), *SBP3a* (**F**), and *SPB4b* (**G**) genes in MT, *SP5G^pen^*, *miR156OE*, *pro*, and *sft* vegetative apices in LD (ZT=4; mean ± SEM; n = 3 samples). **(H-K)** Relative expression of *DST* (**H**), *FA* (**I**), *FUL1* (**J**), and *TM3* (**K**) genes in *sft*, *sft pro*, *sft* *miR156OE*, and *sft SP5G^pen^* vegetative apices in LD (ZT=4; mean ± SEM; n=3 samples). Distinct letters indicate significant differences according to Tukey’s test, *p<0.05*. Each dot represents individual data. MT plants shown in Figure S17A–17C are the same as those used in Figure 10A–10C.

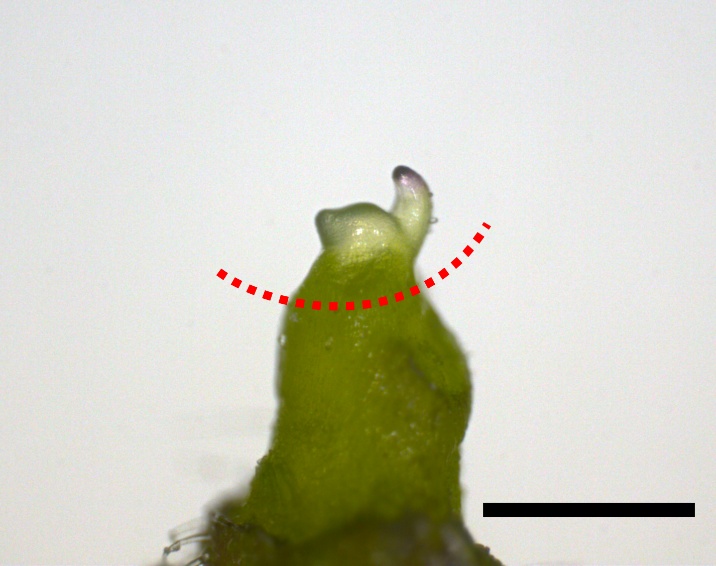

**Figure S18.** **Representative image of a tomato shoot apex harvested for qRT-PCR assays.** The red line indicates where the apices were cut during collection. Bar = 500 µm.

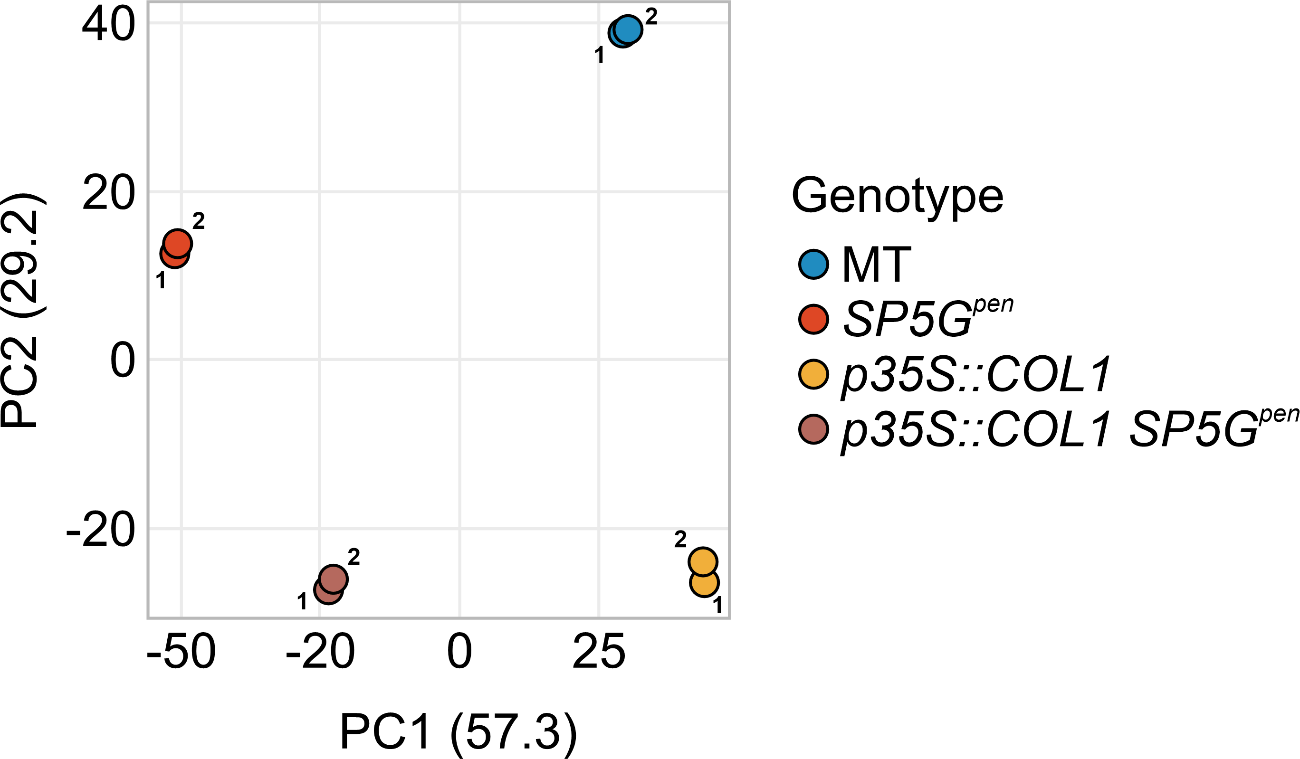

**Figure S19. Principal component analysis (PCA) of transcriptomic data showing the separation of samples according to genotype.** Samples from MT, *SP5G^pen^*, *p35S::SlCOL1*, and *35S::SlCOL1 SP5G^pen^* form distinct clusters, reflecting clear differences in transcriptomic profiles. Biological replicates (1 and 2) cluster closely within each group, indicating high reproducibility.

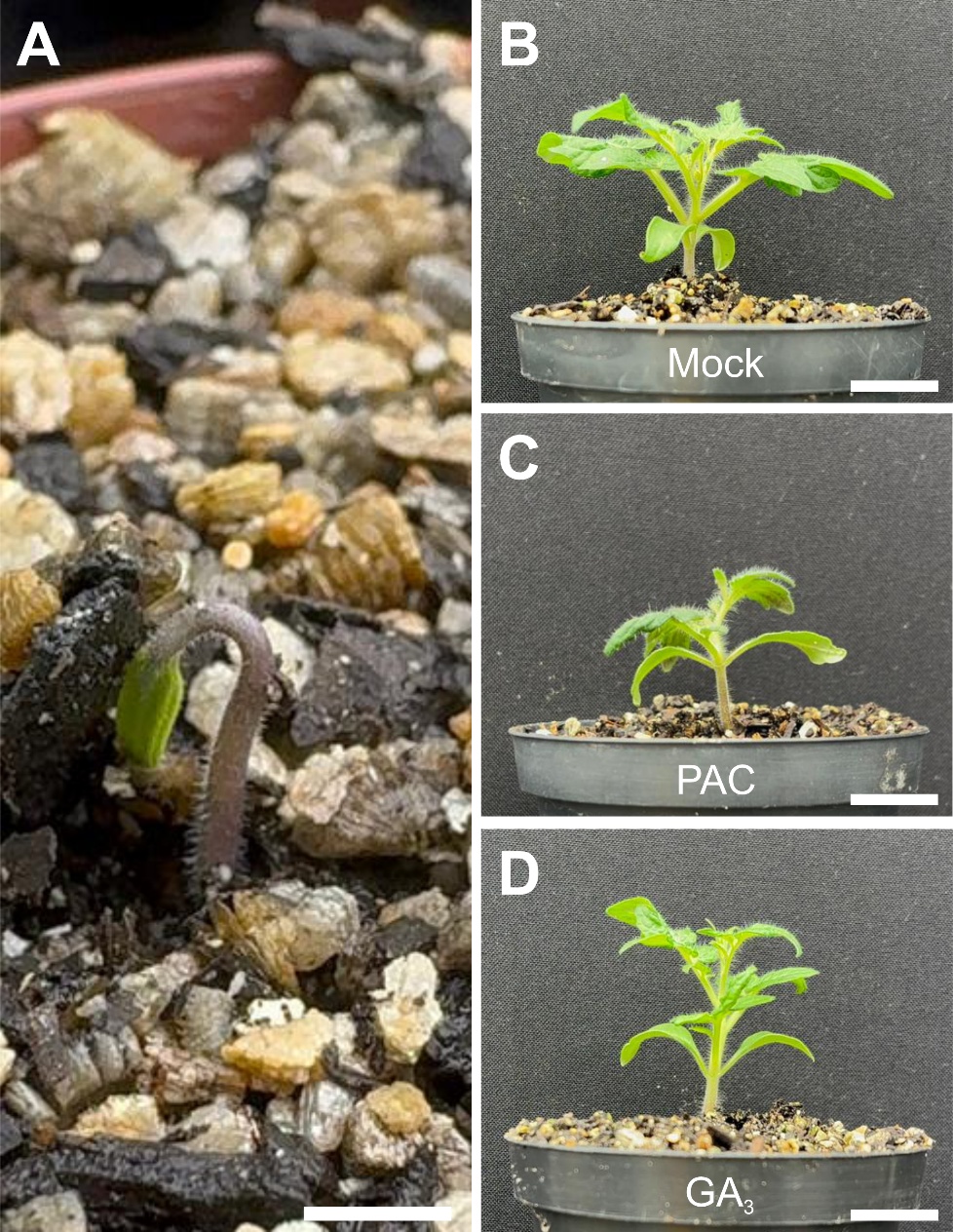

**Figure S20. Representative images of tomato seedlings pre- and post-treatment with GA₃ and PAC. (A)** Seedling at the initial stage of treatment (first day after emergence). Bar = 5 mm **(B–D)** Representative pictures of 10-day-old seedlings of MT after treatment with mock (**B**), PAC (**C**), and GA_3_ (**D**) cultivated in LD (14h/10h). Bar = 5 cm. Treatments were applied by soil drenching.
